# Wheat transcriptome profiling reveals abscisic and gibberellic acid treatments regulate early-stage phytohormone defense signaling, cell wall fortification, and metabolic switches following *Fusarium graminearum*-challenge

**DOI:** 10.1101/2020.09.17.302737

**Authors:** Leann M. Buhrow, Ziying Liu, Dustin Cram, Tanya Sharma, Nora A. Foroud, Youlian Pan, Michele C. Loewen

## Abstract

**Background:** Application of the wheat phytohormones abscisic acid (ABA) or gibberellic acid (GA) affect Fusarium head blight (FHB) disease severity; however, the molecular underpinnings of the elicited phenotypes remain unclear. Herein, the transcriptomic responses of an FHB-susceptible wheat cultivar ‘Fielder’ were characterized upon treatment with ABA, an ABA receptor antagonist (AS6), or GA in the presence or absence of *Fusarium graminearum* (*Fg*) challenge.

**Results:** A total of 30,876 differentially expressed genes (DEGs) where identified in ‘Fielder’ (26,004) and *Fg* (4,872). *Fg* challenge alone resulted in the most substantial wheat DEGs contributing to 57.2% of the total transcriptomic variation. Using a combination of topology overlap and correlation analyses, 9,689 *Fg*-related wheat DEGs were defined. Further enrichment analysis of the top 1% networked wheat DEGs identified critical expression changes within defense responses, cell structural metabolism, molecular transport, and membrane/lipid metabolism. *Fg*-challenged conditions also included the expression of a putative *Fg* ABA-biosynthetic cytochrome P450 and repression of wheat *FUS3* for dysregulating ABA and GA crosstalk. ABA treatment alone elicited 4536 (32%) wheat DEGs common to those of the *Fg*-challenge, and *Fg*+ABA further enhanced 888 (12.5%) of them. These ABA elicited DEGs are involved in defense through both classical and non-classical phytohormone signaling and regulating cell wall structures including polyphenolic metabolism. Conversely, *Fg*+GA opposed 2239 (33%) *Fg*-elicited wheat DEGs, including modulating primary and secondary metabolism, defense responses, and flowering genes. ABA and jointly ABA⍰*Fg*⍰[*Fg*+ABA] treatments repressed, while *Fg*+GA induced an over-representation of wheat DEGs mapping to chromosome 6BL. Finally, compared to *Fg*+ABA, co-application of *Fg*+AS6 did not antagonize ABA biosynthesis or signal but rather elicited antagonistic *Fg* (557) and wheat (11) DEGs responses directly tied to stress responses, phytohormone transport, and FHB.

**Conclusions:** Comparative transcriptomics highlight the effects of wheat phytohormones on individual pathway and global metabolism simultaneously. Application of ABA may reduce FHB severity through misregulating defense mechanisms and cell wall fortification pathways. GA application may alter primary and secondary metabolism, creating a metabolic shift to ultimately reduce FHB severity. By comparing these findings to those previously reported for four additional plant genotypes, an additive model of the wheat-*Fg* interaction is proposed.

## BACKGROUND

Fusarium head blight (FHB), one of the more prevalent diseases of wheat (*Triticum aestivum* L.), is the result of infection of wheat heads by the hemi-biotrophic ascomycetous *Fusarium graminearum* (*Fg*) and related species [1]. FHB remains a significant pathogenic threat to the agricultural industry; *Fg* infection decreases grain yields and deposits mycotoxins [2, 3]. Wheat breeding programs aimed at addressing FHB have resulted in incremental benefits to date [4], while chemical control measures remain limited by host-developmental requirements of the pathogen [5]. Thus, it is imperative that we expand our understanding of these host-pathogen interaction to identify of new sources of resistance and antifungal molecular targets.

The transcriptomic responses of FHB-susceptible and -resistant wheat varieties in response to *Fg* challenge have been investigated extensively over the course of the last decade (reviewed in [6]; and more recently including [7–11]). Together these studies highlighted a breadth of host-responses that include modulation of primary metabolism and photosynthesis, transcriptional and translational regulators, traditional plant defense responses (both phytohormone-associated and pathogenesis-related protein targets), and detoxification genes. Comparative transcriptomics have been applied to *Fg*-challenged genotypes for FHB resistance QTLs 2DL, *Fhb1, Fhb2* and *Qfhs.ifa-5A* with outcomes emphasizing increased basal defenses, enhanced classical phytohormone defense signaling, and antagonism of pathogen-mediated modulation of phytohormone pathways [12–16]. The role of the classical defense phytohormones, salicylic acid (SA), jasmonic acid (JA), and ethylene (ET) in the wheat defense response to *Fg*-challenge has been extensively described with a consensus model of early biotrophic SA followed by later stage necrotrophic JA/ET responses [6, 8, 9]. Nonetheless, the role of ET remains in question, with conflicting reports highlighting both mediation of resistance [3] as well as susceptibility [17] in early and late responses [9].

In agreement with independent research groups investigating other wheat varieties and FHB disease stages [9,18,19], our previous report described the drastic alternation of phytohormone profiles in the FHB-susceptible *T. aestivum* cultivar ‘Fielder’ when challenged with *Fg* [20]. Two such FHB-regulated phytohormones, when co-applied with pathogen challenge, modulated disease severity and spread where abscisic acid (ABA) promoted infection and gibberellic acid (GA) reduced infection [19, 20]. Comparative analysis of the *Fg* transcriptomic responses to the presence of these phytohormones, measured at 24 h post treatment (hpt), emphasized promotion of early-infection genes in the presence of ABA and reduced nitrogen metabolism in the presence of GA [20]. It has also been established that *Fusarium* spp. can themselves produce ABA [19], GA [21], auxin (indole acetic acid; IAA [22]), and cytokinins (CK; [23]), while also encoding both 1-aminocyclopropane carboxylic acid (ACC) synthases and deaminases potentially involved in ET biosynthesis [24]. Therefore, although phytohormones may be traditionally thought to serve as plant host signaling and defense molecules, it is unclear how *Fusarium* spp. may dysregulate phytohormone metabolism to establish or promote infection.

In this light, the present study aims to identify and characterize ‘Fielder’ host transcriptomic responses to *Fg* challenge, ABA or GA treatment, and the combined effects of both the pathogen and one of the two phytohormones. Additionally, this analysis considers the effects on the host and pathogen when treated with a co-applied antagonist of ABA receptors, AS6 [25], which was previously reported to have an insignificant phenotypic effect on FHB late-stage disease severity [20]. Notably to ensure the most up-to-date interpretations, all RNA-seq data was processed using the recently completed version wheat genome sequences, IWGSC RefSeq v1.0 [26]. Finally, we compare transcriptomic responses of five wheat genotypes upon *Fusarium* challenge to further contribute to a consensus model of these plant-pathogen interactions. We discuss how transcriptomic changes elicited by ABA or GA impact such a consensus model and may contribute to the modulation of FHB symptoms previously reported in Qi et al. [19] and Buhrow et al. [20].

## RESULTS

### Transcriptome overview

FHB-susceptible wheat cultivar ‘Fielder’ spikes were treated with ABA (condition: ABA), GA (condition: GA), or *Fg* (condition: *Fg*) alone. Additional ‘Fielder’ spikes were treated with a combination of *Fg* and ABA (condition: *Fg*+ABA), GA (condition: *Fg*+GA) or an ABA receptor antagonist (condition: *Fg*+AS6). Twenty-four hours after the treatments, RNA was extracted from wheat spikelets in each treatment group (RNA Integrity Number (RIN) = 8.9 ± 0.38) and sequenced with an average of 27 ± 7.7 million pair-end RNA-seq reads per sample **(Additional File 1 Tab ‘S1’).** A subset of this RNA-seq data was previously reported on *F. graminearum* differential gene expression (DEG) upon phytohormone treatment [20]. In the present study, RNA-seq reads were remapped to a combination of the recently completed version of wheat genome (IWGSC RefSeq v1.0; [26]) and *Fusarium graminearum* (str. PH-1) resulting in 97 % of the transcripts being successfully mapped to the two reference genomes **(Additional file 1, Tab ‘S1’).** *Fg*-challenge and phytohormone application resulted in consistent and distinguishable changes to the ‘Fielder’ transcriptome **(Figure 1).** After normalization of the read counts, a total of 30,876 differentially expressed genes (DEGs) were identified based on the criteria specified in the method section, including 4,872 from *F. graminearum* and 26,004 from wheat **(Table 1; Additional Files 2 & 3).**

**Figure 1:**
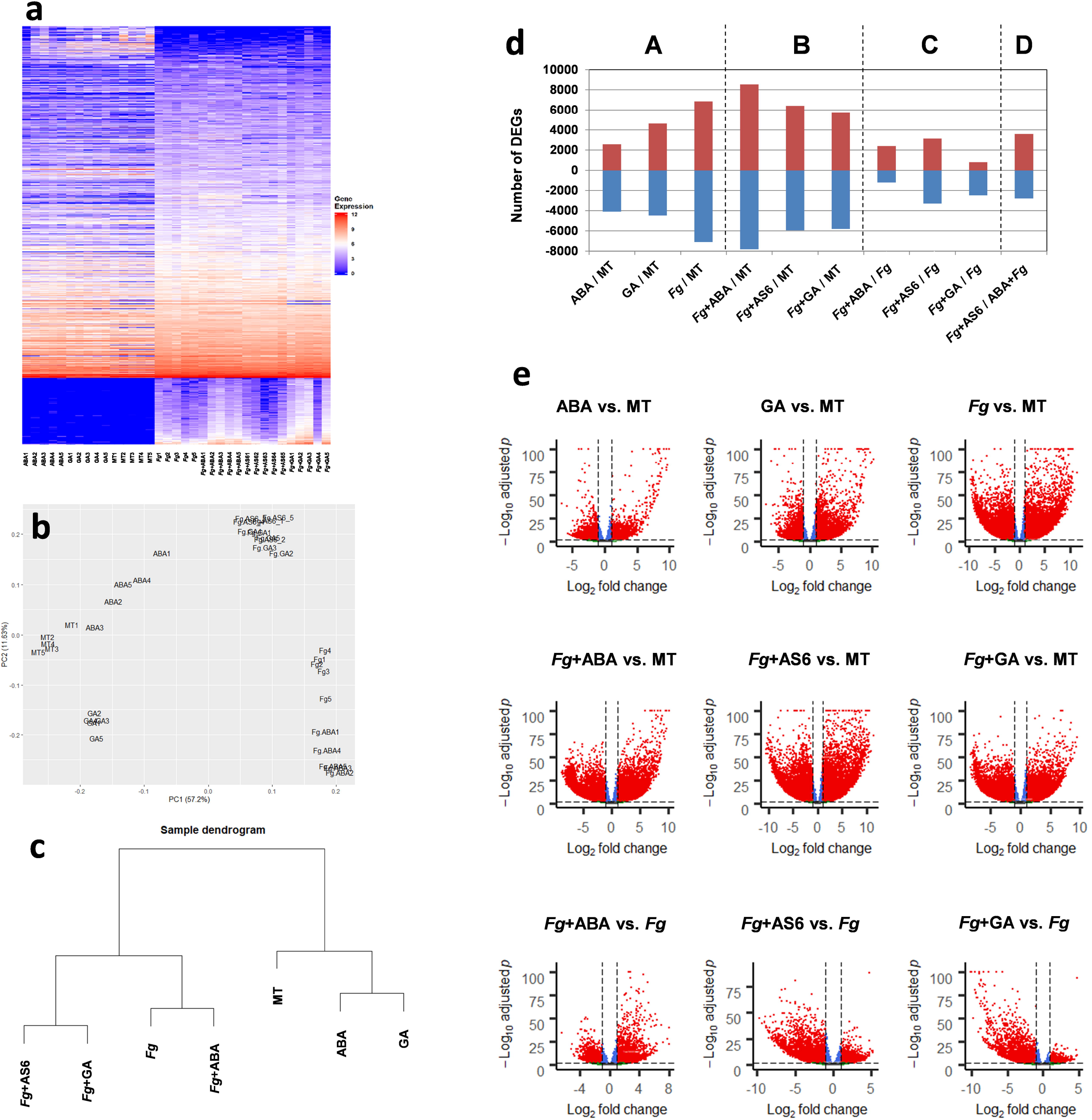
Transcriptome overview. Gene expression levels were log2 transformed transcript counts, **a)** heatmap of 30,876 DEGs, the top portion is for wheat genes and the bottom portion for *Fg* genes; **b)** PCA of wheat DEGs; **c)** sample dendrogram of wheat DEGs; **d)** wheat DEG distribution over the 10 pairwise comparisons, A: “alone” conditions compared with MT, B: co-applications compared with MT, C: co-applications compared with *Fg,* D: comparison between two co-applications; **e)** volcano plots of wheat DEGs of the first nine pairwise comparisons (sub-panel A, B, C of panel d), adjusted *p* values smaller than 10^−100^ were displayed as 10^−100^.

**Table 1.** Number of DEGs arising from each pairwise comparison.

| DEG Population | Regulation | Condition |  |  |  |  |  |  |  |  |  |
| --- | --- | --- | --- | --- | --- | --- | --- | --- | --- | --- | --- |
|  |  | ABA /MT | GA /MT | *Fg* /MT | *Fg*+ABA /MT | *Fg*+AS6 /MT | *Fg*+GA /MT | *Fg*+ABA /*Fg* | *Fg*+GA /*Fg* | *Fg*+AS6 /*Fg* | *Fg*+AS6 /*Fg*+ABA |
| Total | Up | 2566 | 4623 | 8691 | 11772 | 7668 | 10560 | 2477 | 1703 | 3163 | 3602 |
|  | Down | 4098 | 4473 | 7100 | 7810 | 5962 | 5798 | 1213 | 2472 | 3325 | 3330 |
| Wheat | Up | 2566 | 4623 | 6855 | 8529 | 6410 | 5743 | 2398 | 806 | 3146 | 3602 |
|  | Down | 4098 | 4473 | 7100 | 7810 | 5962 | 5798 | 1213 | 2471 | 3299 | 2773 |
| *Fg* | Up | 0 | 0 | 1836 | 3243 | 1258 | 4817 | 79 | 897 | 17 | 0 |
|  | Down | 0 | 0 | 0 | 0 | 0 | 0 | 0 | 1 | 26 | 557 |

Toward assessing the relative effects of the different treatments, differential expression feature extraction (DEFE; [27]) analysis was performed. Among the 729 (3^6^) theoretically possible ‘M’ DEFE-patterns for the six treatments as they compared with mock water treatment (MT), 266 had one or more genes. A list of the 30 highly populated DEFE patterns collectively containing 17,170 (66 %) of the wheat DEGs and 4,839 **(99 %)** of the *Fg* DEGs **(Table 2).** The frequency distribution in other **DEFE** patterns is available in **Additional Files 2 & 3 Tab ‘DEFE_stats’**. At a high level, the DEFE analysis was broadly consistent with the sheer impact of the pathogen on wheat gene expression regardless of co-applied treatments, as well as the independent effects of the phytohormones ABA and GA in the absence of the pathogen. Observations of the patterns containing the highest number of wheat DEGs included M001100 (1,736 DEGs): up-regulated by the pathogen with or without co-application of ABA, followed by M202222 (1,641 DEGs): down-regulated by pathogen with or without co-application of any of ABA, GA or AS6, and by ABA-alone (no pathogen). These were followed closely by up-regulated (M001111; 1,593 DEGs) or down-regulated (M002222; 1,181 DEGs) by *Fg,* with or without co-application of ABA, GA or AS6, as well as up-regulated (M0100000; 1306 DEGs) and down-regulated (M020000; 1,163 DEGs) by GA with no pathogen infection. Similarly, *Fg* DEGs were highly enriched with four patterns: (1) M000001 (1,589, DEGs): up-regulated by co application of GA and the pathogen; (2) M001111 (1,165 DEGs): up-regulated by pathogen infection, with or without co-application of ABA, GA or AS6; (3) M000101 (1,315 DEGs): up-regulated by co-application of the pathogen with either ABA or GA; and (4) M001101 (647 DEGs): up-regulated by the pathogen, with or without co-application of ABA or GA. These four patterns collectively accounted for 97 % of *Fg* DEGs **(Table 1, Additional File 3).**

**Table 2.**
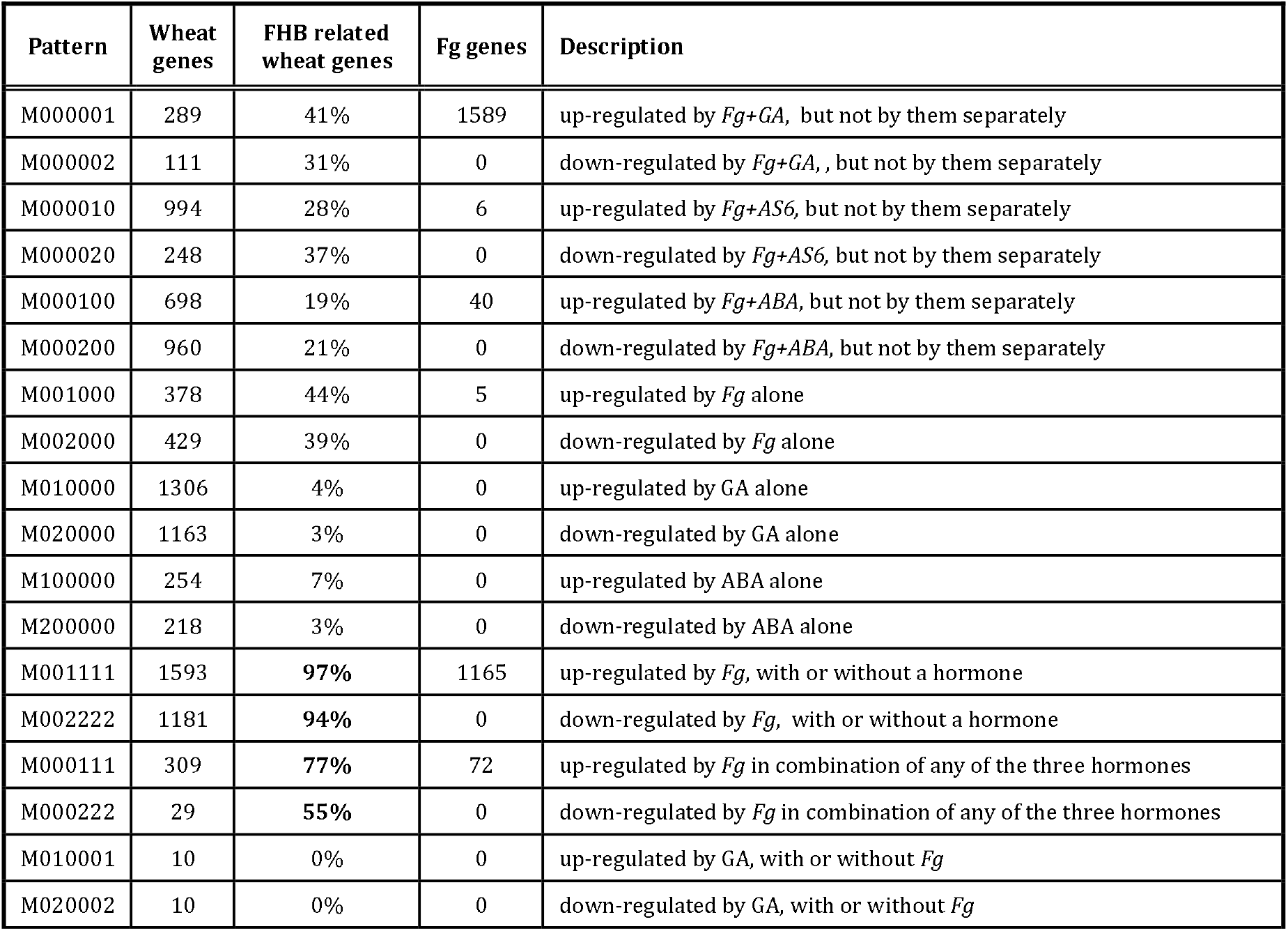
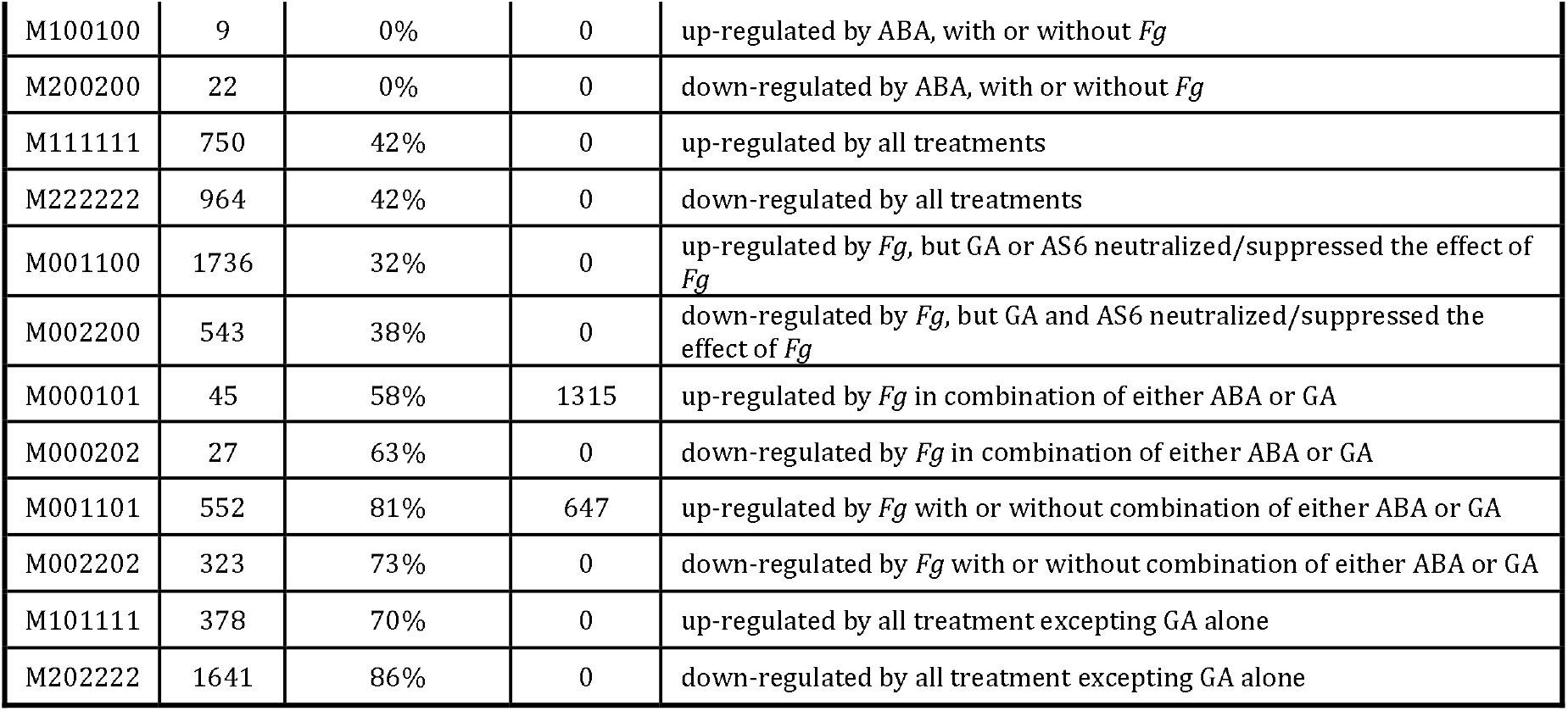
Number of genes in highly informative <u>D</u>ifferential <u>E</u>xpression <u>F</u>eature <u>E</u>xtraction (DEFE) patterns M(ABA/MT, GA/MT, *Fg*/MT, *Fg*+ABA/MT, *Fg*+AS6/MT, *Fg*+GA/MT)

### Wheat DEGs highly correlated with *Fg* challenge

Wheat transcriptomic responses to pathogen challenge can be clearly distinguished based on the presence or absence of *Fg* as evidenced by the principle component analysis of DEGs (57.2% of variance; **Figure 1b).** Through topology overlap analysis using WGCNA [28], 58 clusters were identified and 17 were significantly correlated (p < 5 x 10^-2^) to *Fg* treatment, with or without co-application of a phytohormone or inhibitor **(Additional File 2 Tab ‘WheatDEGs’ @Col-M,N & Tab ‘moduleTraitCor’).** These 17 clusters collectively contained 12,138 DEGs. This finding was further validated by a separate correlation analysis which revealed 10,609 individual *Fg*-correlated DEGs, of which, 9,689 (91 %) were also in the 17 clusters, and thus considered as ‘*Fg*-related genes’ **(Additional File 2 Tab ‘WheatDEGs’ @Col-O).** The majority (83 %) of these 9,689 *Fg*-related ‘Fielder’ genes fall into the most informative ‘M’ DEFE patterns listed in **Table 2.**

Gene association network analysis was performed based on the correlation topology overlap detected among the DEGs using the WGCNA R package [28]. The top 1 % of the topology overlap matrix was considered in the entire network consisting of 10,373 wheat DEGs, connected by 3,370,764 edges. Of these, 8,349 (80 %) were *Fg*-related genes. The top 2,621 genes having connection degrees of 1,000 or higher were considered key hub genes (highlighted in **Additional File 2 Tab ‘WheatDEGs’@Col-P).** An enrichment index analysis of gene groups was performed on the network genes as described in Pan et al., [8]. Among the highly enriched group of genes in the network were alkaline shock protein 23, clathrin assembly family protein, D-glycerate dehydrogenase / hydroxypyruvate reductase, photosystem II 22 kDa, chloroplastic, PISTILLATA-like MADS-box transcription factor, fatty acid hydroxylase, fimbrin-like protein 2, pollen allergen, glyoxal oxidase, jasmonate ZIM domain proteins, C2 domain-containing protein, yellow stripe-like transporters, and pectinesterase **(Additional File 4).** The B3 domain-containing transcription factor *FUS3* gene (TraesCS3D01G249100) was among the key hub genes, with 2,281 immediate neighborhood genes in the network and was significantly down-regulated by all conditions that include *Fg*-treatment (adj p < 2 x 10^-14^) **(Additional File 5: Figure S1).** Interestingly, these conditions that include *Fg*-treatment also express a putative fungal ABA biosynthetic cytochrome P450 homolog (FGRAMPH1-01T26277, **Additional File 5: Figure S2B).**

### ABA and GA treatments alone elicit diverse metabolic changes to the wheat transcriptome

The impact of ABA and GA application alone was considered with DEGs compared to MT. Neither phytohormone treatment elicited wheat responses as strong as *Fg* challenge **(Figures 1a & b; Table 1).** ABA treatment down-regulated nearly twice as many wheat genes as it up-regulated, while GA modified approximately equal numbers of genes up and down **(Table 1, Figures 1d & e),** consistent with the trends recently reported on the FHB-susceptible wheat cultivar ‘Roblin’ [11]. ABA treatment most notably led to downregulation of 3079 (43 %) wheat DEGs that were down-regulated by *Fg* and upregulation of 1457 (21 %) wheat DEGs that were up-regulated by *Fg* **(Figure 2).** Furthermore, ABA elicited few opposing effects when compared to the *Fg* condition, downregulating 86 and up-regulating 62 wheat DEGs. A GO enrichment analysis of the opposing 86 down-regulated wheat DEGs highlights aromatic compound biosynthesis (10 hits, p < 5 x 10^−4^) and cellular nitrogen compound biosynthesis (10 hits, p < 5 x 10^−3^) **(Additional File 6 Tab ‘FU2’).** While GA treatment elicited 20 – 25 % of wheat DEGs in the same directionality as the *Fg* condition, it is decidedly more notable that GA down-regulated 204 (3 %) DEGs that were up-regulated by *Fg* and conversely up-regulating 286 (4 %) DEGs that were down-regulated by *Fg* **(Figure 2).** Gene ontology (GO) enrichment analysis of the down-regulated 204 genes yields 70 hits (p < 7 x 10^−3^) with either hydrolase (39 hits, p < 10^−5^, including glucosyl and ester) or oxidoreductase activities (23 hits, p < 7 x 10^−3^), heme binding (12 hits; p < 3 x 10^−3^), carbohydrate metabolite processes (17 hits, p < 10^−4^) and catabolic processes (12 hits, p < 6 x 10^−3^) **(Additional File 6 Tab ‘FU4’).** Interestingly with respect to the up-regulated 286 DEGs, 85 hits were associated with primary metabolic processes (p < 3 x 10^−2^) and 90 hits for organic substance metabolic processes (p < 2 x 10^−2^) including 22 hits under lipid metabolic process (p < 2 x 10^−5^) and 20 hits under carbohydrate metabolic process (p < 10^−3^).

**Figure 2:**
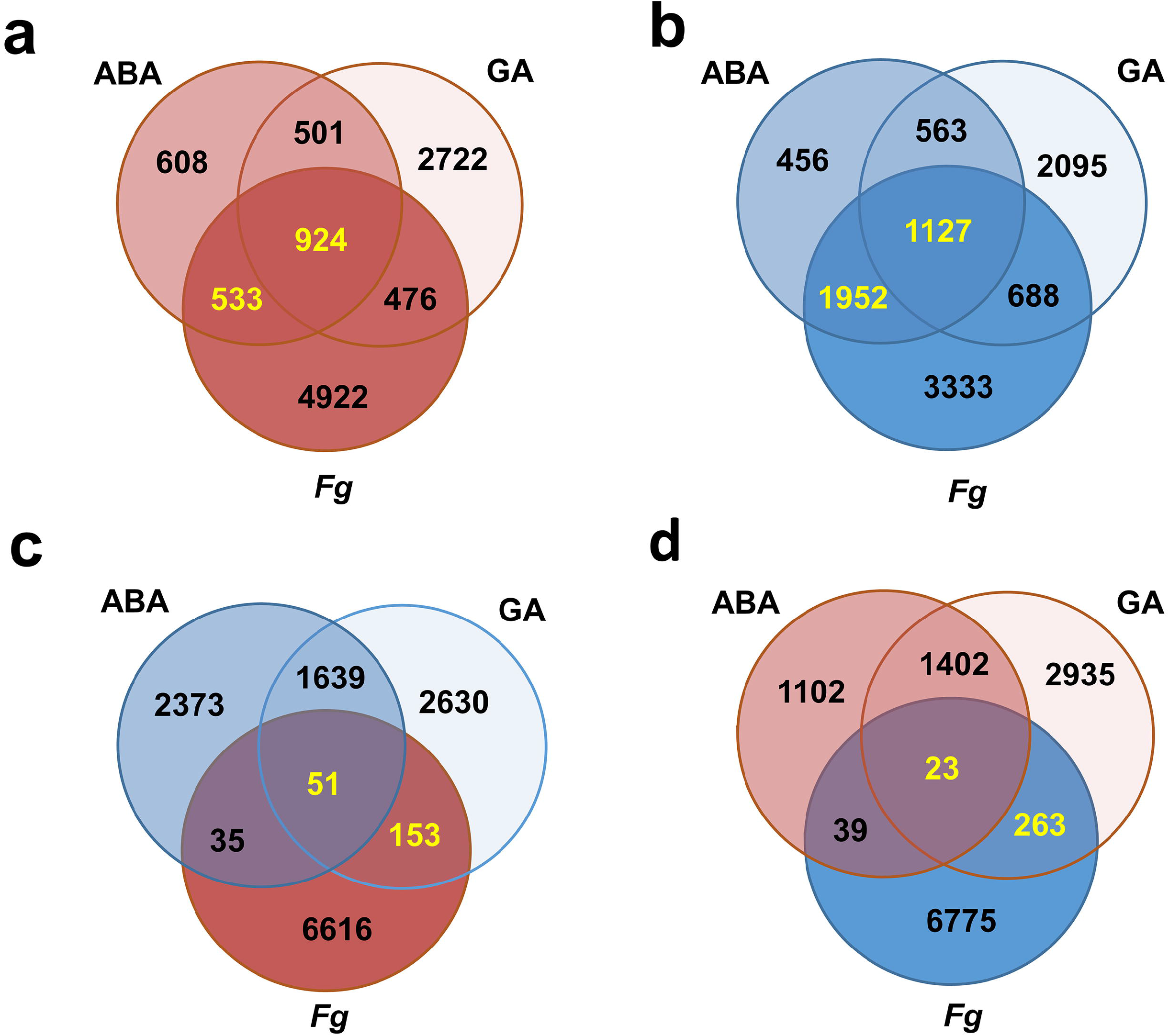
Share of *Fg* alone up- or down-regulated genes that are up- and down-regulated under ABA and GA alone conditions, with all DEGs calculated compared to mock treated (MT). In all cases, up-regulated genes are highlighted in shades of red, down-regulated genes are highlighted in shades of blue. DEGs called in text are yellow. **a)** Up-regulated DEGS. **b)** Down-regulated DEGS. Opposing DEGs: **c)** up-regulated by *Fg* but down-regulated by ABA and GA; **d)** down-regulated by *Fg* but up-regulated by ABA and GA.

### ABA or GA co-application respectively with *Fg* enhanced and suppressed wheat DEGs elicited by *Fg* alone

The wheat transcriptomic changes elicited by ABA (*Fg*+ABA) and GA (*Fg*+GA) co-application in the presence of *Fg* pathogen challenge were collectively analyzed as compared to MT thus allowing a direct comparison of gene regulation events mediated by phytohormones alone to those arising from the condition where all three components of the ‘plant-pathogen-phytohormone’ interaction were present together. In the case of responses common to the *Fg, Fg*+ABA and *Fg*+GA treatments (*Fg*⍰[*Fg*+ABA]⍰[*Fg*+GA]), these conditions jointly enhanced 3751 (55 %) wheat DEGs that were up-regulated by *Fg* alone and suppressed 4696 (66 %) that were DEGs down-regulated by *Fg* alone (highlighted in **Additional File 5: Figures S3A & B).** The co-applications also elicited opposing wheat DEGs where *Fg*+ABA enhanced 63 and suppressed four DEGs, and *Fg*+GA enhanced two and suppressed five DEGs that were down-regulated and up-regulated by *Fg* alone (highlighted in **Additional File 5: Figures S3C & D).** These changes, along with non-overlapping DEGs, manifested in an increase upon *Fg*+ABA and slight reduction upon *Fg*+GA treatments in the overall DEG counts when compared to *Fg* alone **(Figure 1d; Table 1).** When further considering DEGs elicited by the phytohormone, pathogen, and co-applied conditions, the overlap in elicited wheat transcriptomic changes was over 90 % DEGs common in the case of ABA (ABA⍰*Fg*⍰[*Fg*+ABA]) and over 80 % DEGs common in the case of GA (GA⍰*Fg*⍰[*Fg*+GA]; **Additional File 5: Figure S4).**

To more specifically assess enhancement and suppression of the host response to *Fg* by the co-applied phytohormones, a second DEFE analysis was carried out as compared to *Fg* challenge **(Table 3; Additional File 2 Tab ‘DEFE_stat’).** The *Fg*+ABA co-application significantly enhanced 316 up-regulated and 572 down-regulated *Fg*-elicited wheat DEGs **(Figures 3a & 3d)** while also inhibiting 142 up-regulated and 279 down-regulated *Fg*-elicited wheat DEGs **(Figures 3c & 3b).** The *Fg*+GA co-application significantly enhanced 152 up-regulated and 349 down-regulated *Fg*-elicited wheat DEGs (highlighted in **Figures 4a & 4d)** while also inhibiting 2,232 (33 %) up-regulated and 7 down-regulated *Fg*-elicited wheat DEGs (highlighted in **Figures 4c & 4b).** These findings clearly demonstrated the additive effects of *Fg*+ABA and the decremental effects of *Fg*+GA on the ‘Fielder’ transcriptome challenged with *Fg* and are consistent with these observations **(Figure 1e),** where ABA co-application predominantly enhances, and GA predominantly represses, the wheat transcriptome.

**Table 3.** Number of genes with highly informative DEFE patterns on enhancement or suppression of a hormone in co-application with *Fg*: F(*Fg*+ABA_vs_*Fg, Fg*+AS6_vs_*Fg, Fg*+GA_vs_*Fg*)

| Pattern | wheat | *Fg* | Description |
| --- | --- | --- | --- |
| F001 | 212 | 856 | Enhanced by GA |
| F002 | 300 | 1 | Suppressed by GA |
| F010 | 2046 | 6 | Enhanced by AS6 |
| F020 | 1036 | 24 | Suppressed by AS6 |
| F100 | 1639 | 39 | Enhanced by ABA |
| F200 | 1038 | 0 | Suppressed by ABA |
| F011 | 381 | 2 | Enhanced by both AS6 and GA, but not by ABA |
| F022 | 2134 | 0 | Suppressed by both AS6 and GA, but not by ABA |
| F101 | 63 | 31 | Enhanced by both ABA and GA, but not by AS6 |
| F202 | 14 | 0 | Suppressed by both ABA and GA, but not by AS6 |
| F110 | 548 | 3 | Enhanced by both ABA and AS6, but not GA |
| F220 | 93 | 0 | Suppressed by both ABA and AS6, but not by GA |
| F111 | 123 | 6 | Enhanced by all three hormones |
| F222 | 13 | 0 | Suppressed by all three hormones |
| F012 | 1 | 0 | Enhanced by AS6, but suppressed by GA |
| F021 | 1 | 2 | Suppressed by AS6, but enhanced by GA |
| F102 | 3 | 0 | Enhanced by ABA, but suppressed by GA |
| F201 | 8 | 0 | Suppressed by both ABA, but enhanced by GA |
| F120 | 16 | 0 | Enhanced by ABA, but suppressed by AS6 |
| F210 | 29 | 0 | Suppressed by both ABA, but enhanced by AS6 |
| F122 | 6 | 0 | Enhanced by ABA, but suppressed by both AS6 and GA |
| F211 | 18 | 0 | Suppressed by ABA, but enhanced by both AS6 and GA |

**Figure 3.**
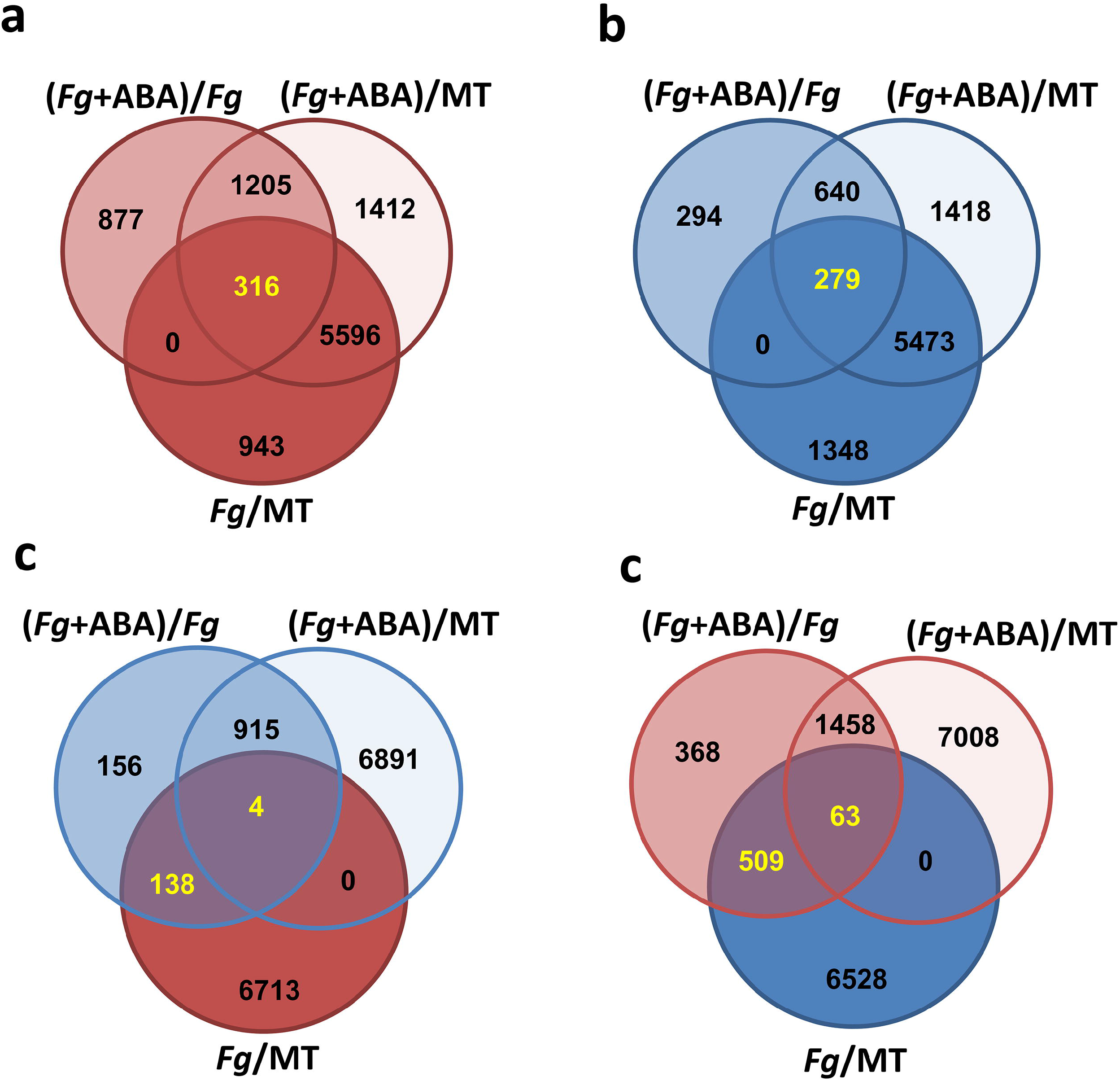
Comparison of gene regulation elicited by ABA in the presence of *Fg,* with DEGs calculated compared to mock treated (MT) as well as to *Fg.* In all cases, up-regulated genes are highlighted in shades of red, and down-regulated genes are highlighted in shades of blue. DEGs called in text are yellow. **a)** Distribution of all up-regulated genes by *Fg*+ABA as compared to *Fg,* or *Fg*+ABA and *Fg*-alone compared to MT. **b)** Distribution of all down-regulated genes by *Fg*+ABA as compared to *Fg,* or *Fg*+ABA and *Fg*-alone compared to MT. **c)** Distribution of up-regulated genes by *Fg*-alone, but down-regulated by *Fg*+ABA as they compared to MT or to *Fg.* **d)** Distribution of all down-regulated genes by *Fg*-alone, but up-regulated by *Fg*+ABA as they compared to MT or to *Fg.*

**Figure 4.**
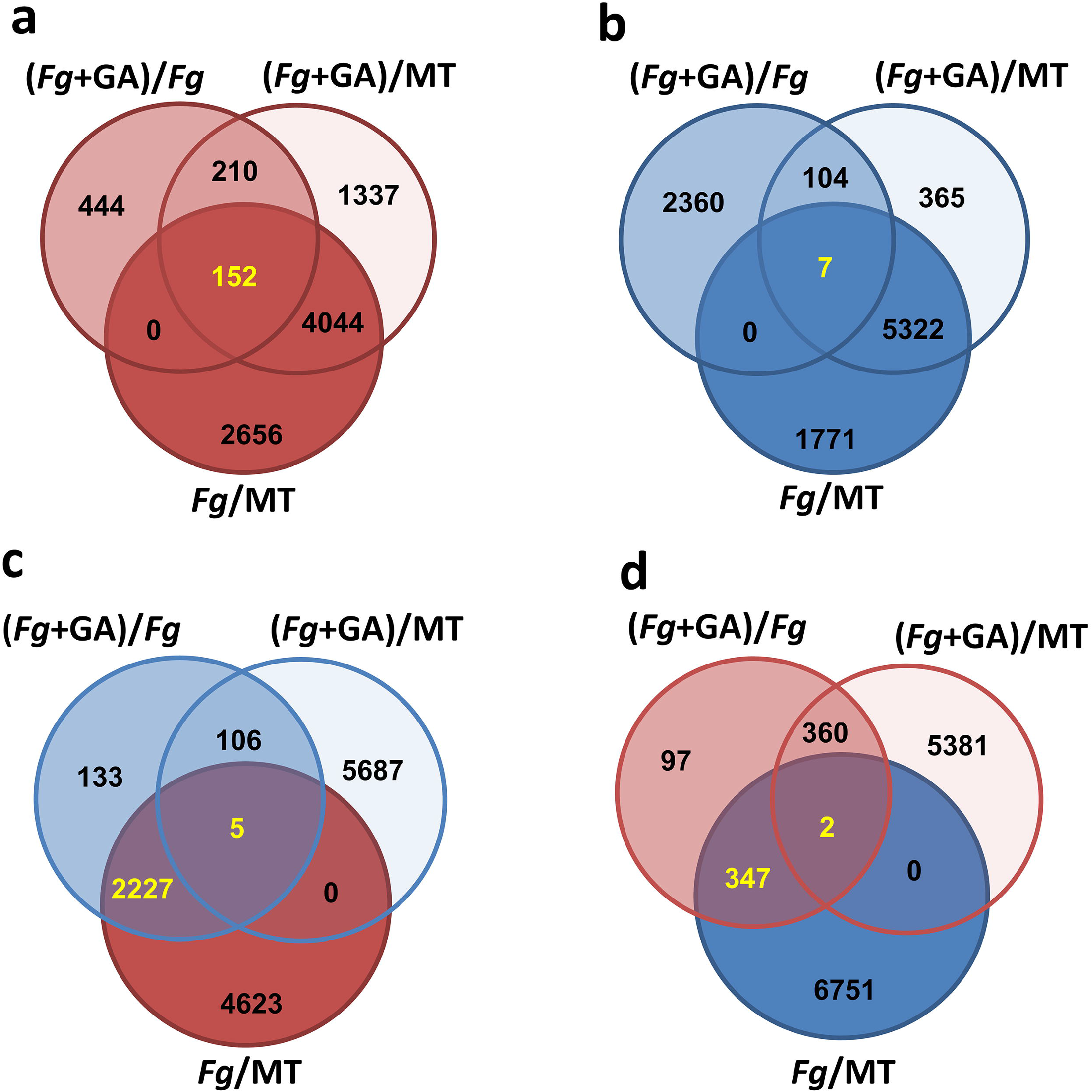
Comparison of gene regulation elicited by GA in the presence of *Fg,* with DEGs calculated compared to mock treated (MT) as well as *Fg*. In all cases, up-regulated genes are highlighted in shades of red, and down-regulated genes are highlighted in shades of blue. DEGs called in text are yellow. **a)** Distribution of all up-regulated genes by *Fg*+GA as compared to *Fg,* or *Fg*+GA and *Fg*-alone compared to MT. **b)** Distribution of all down-regulated genes by *Fg*+GA as compared to *Fg,* or *Fg*+GA and *Fg*-alone compared to MT. **c)** Distribution of up-regulated genes by *Fg*-alone, but down-regulated by *Fg*+GA as they compared to MT or to *Fg.* **d)** Distribution of all down-regulated genes by *Fg*-alone, but up-regulated by *Fg*+GA as they compared to MT or to *Fg.*

### Application of ABA and GA elicit wheat DEGs mapped to chromosome 6BL consistent with their modulation of later-stage FHB phenotype

To understand if the modulation of FHB spread by the co-application of ABA or GA may be tied to a characterized FHB-resistance quantitative trait loci (QTL), DEGs from all treatments were mapped and binned based on chromosomal arm. As expected, DEGs mapped in a bimodal distribution where greater counts were elicited with the MT comparator than *Fg* challenge **(Figures 5a & 5b).** Density as a function of physical length [26] was relatively monodisperse with 26.1 ± 6.4 (average ± one standard deviation) up-regulated DEGs and 25.9 ± 6.3 down-regulated DEGs calculated per Mb, excluding Chr5AS for which fewer induced DEGs were mapped **(Additional File 5: Figure S5).** Additionally, DEGs appear to map uniformly between the A, B, and D genomes with the exceptions of Chr 4AL and 6BL having additional induced DEGs, Chr 5AS and 7BS having fewer induced DEGs, Chr 4AL having additional repressed DEGS, and Chr 7BS having fewer repressed DEGs when compared to their homoeologous chromosome arms **(Figures 5a & 5b).** Interestingly, when divided based on treatment, the greatest mapping in both induced and repressed DEGs is attributed to the long arms of Chr 5, while consistently the fewest mapped DEGs are attributed the short arm of Chr5 **(Additional File 5: Figure S5).** Chr6BL also appears to exhibit differential trends with an over-representation of induced DEGs in ABA and ABA⍰*Fg*⍰[*Fg*+ABA] treatment groups, while also having an over-representation of repressed DEGs upon *Fg*+GA treatment. As these early-stage results are in agreement with the late-stage FHB phenotypes elicited by these treatments, it is possible that the genetic underpinnings may be partially attributed to genes located on Chr 6BL. In wheat and barley, this chromosome arm encodes ABA and GA opposingly-regulated α-amylase genes [29], an ABA 8’-hydroxylase important for ABA catabolism and seed germination [30], and ABA responsive yet-uncharacterized gene(s) that are involved in dormancy [31].

**Figure 5:**
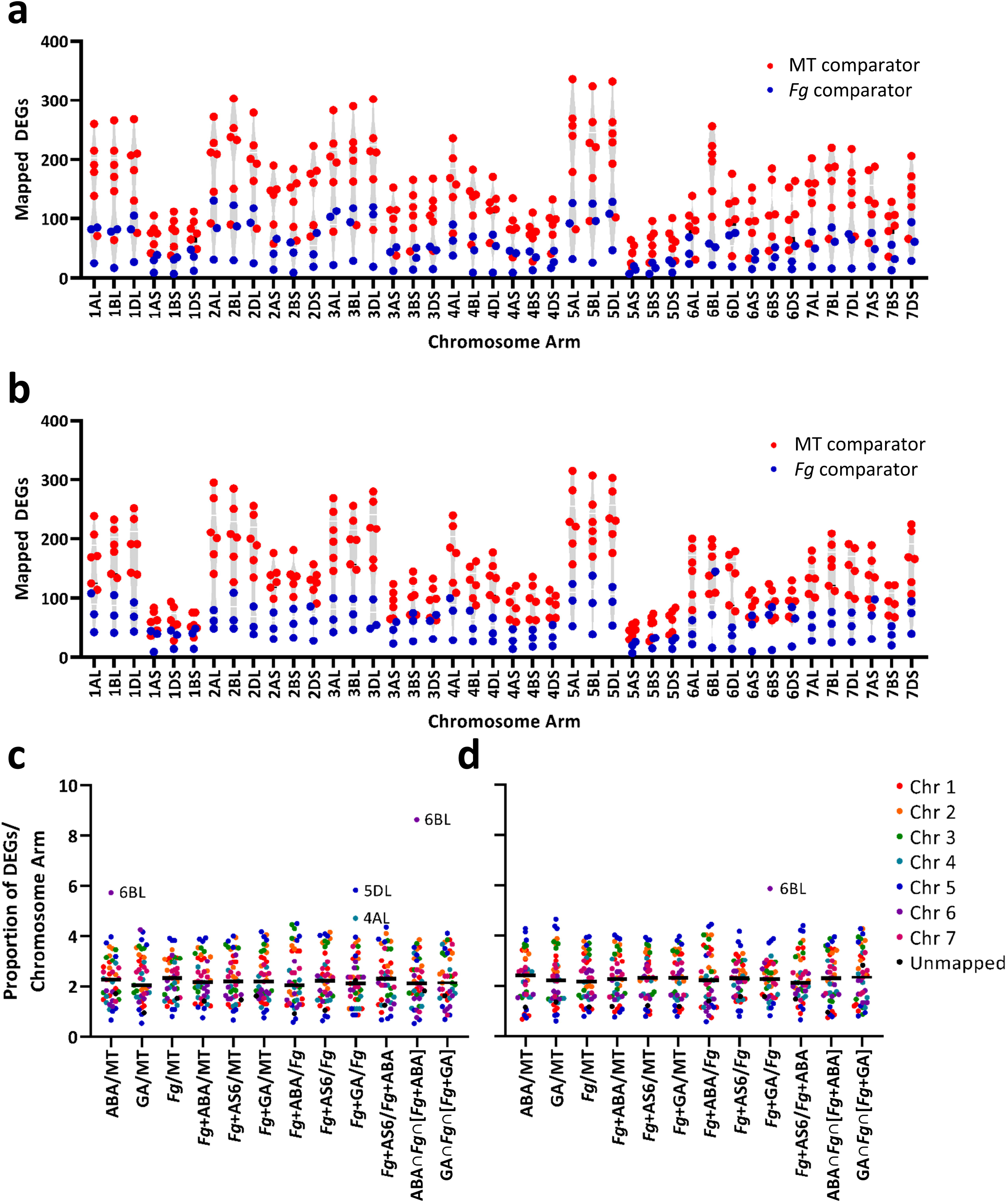
Chromosomal mapping of wheat DEGs elicited by *Fg* and phytohormone treatments. Mapped DEGs as a function of chromosome arm were subdivided into **a)** up-regulated and b) down-regulated DEGs with data points representing individual treatments of *Fg,* phytohormone, or the combination as compared to MT (6 comparisons in red) or *Fg* (3 comparisons in blue). The proportion of DEGs **c)** up-regulated or **d)** down-regulated was also calculated as DEGs per chromosome arm compared to total and was represented as a function of individual treatment and further parsed by chromosome.

### Application of ABA receptor antagonist, AS6, elicits few *Fg* DEGs but recapitulates the wheat DEGs induced by ABA or reduced by GA co-application

With respect to the *Fg* genes, both *Fg*+ABA and *Fg*+GA treatments up-regulated DEGs **(Table 1)** while *Fg*+AS6 elicited a mixed effect on the fungal transcriptome **(Additional File 5: Figure S2A,).** For the *Fg*+AS6 treatment, GO enrichment analysis of the 26 repressed DEGs categorized 11 as putatively having oxidoreductase activity (p < 10^−4^, **Additional File 6 Tab ‘F1’, Additional File 7);** while the 17 enhanced DEGs included three genes have hydrolase activity, hydrolyzing O-glycosyl compounds (p < 4 x 10^−3^, **Additional File 6 Tab ‘F2’).** Interestingly, when this condition was compared to *Fg*+ABA, the presence of AS6 analog repressed 557 DEGs up-regulated by *Fg*+ABA, but had no opposite effect **(Table 1),** supporting the role of this putative antagonist functionally eliciting opposing transcriptomic responses on the pathogen.

On the side of wheat host transcriptomic responses, *Fg*+ABA largely elicited additive while *Fg*+GA largely elicited decremental DEGs when compared to *Fg* challenge alone (refer to previous sections). The *Fg*+AS6 condition was more nuanced as it enhanced the expression of 604 (8.8 %) up-regulated and **904** (13 %) down-regulated wheat DEGs elicited by *Fg* alone **(Additional File 5: Figures S6A & D),** while also inhibiting 2,685 (39 %) up-regulated and 47 (0.66 %) down-regulated wheat DEGs elicited by *Fg* alone **(Additional File 5: Figures S6C & B).** Additionally, the co-application of *Fg*+ABA or *Fg*+AS6 had strong enhancement and suppression **(Table 3,** DEFE patterns F100, F010, F200, F020) effects respectively on gene regulation compared to *Fg*-alone, while co-applied *Fg*+AS6 or *Fg*+GA jointly suppressed 2,134 genes (F022, the highest number of DEGs in F series DEFE analyses, **Table 3**). In summary, *Fg*+AS6 suppressed genes had 86 % overlap with those arising from the *Fg*+GA suppression condition **(Figures 1e & 6c),** while conversely, *Fg*+AS6 enhanced genes had a 23 % overlap with *Fg*+ABA enhancement **(Table 3** DEFE pattern F110). The high repressive effects of *Fg*+AS6 were consistent with the magnitude of differential expression **(Figure 1e);** while the union of both *Fg*+GA and *Fg*+AS6 suppressed wheat DEGs supports the general similarity observed between these two conditions as revealed in the PCA **(Figure 1b).**

**Figure 6.**
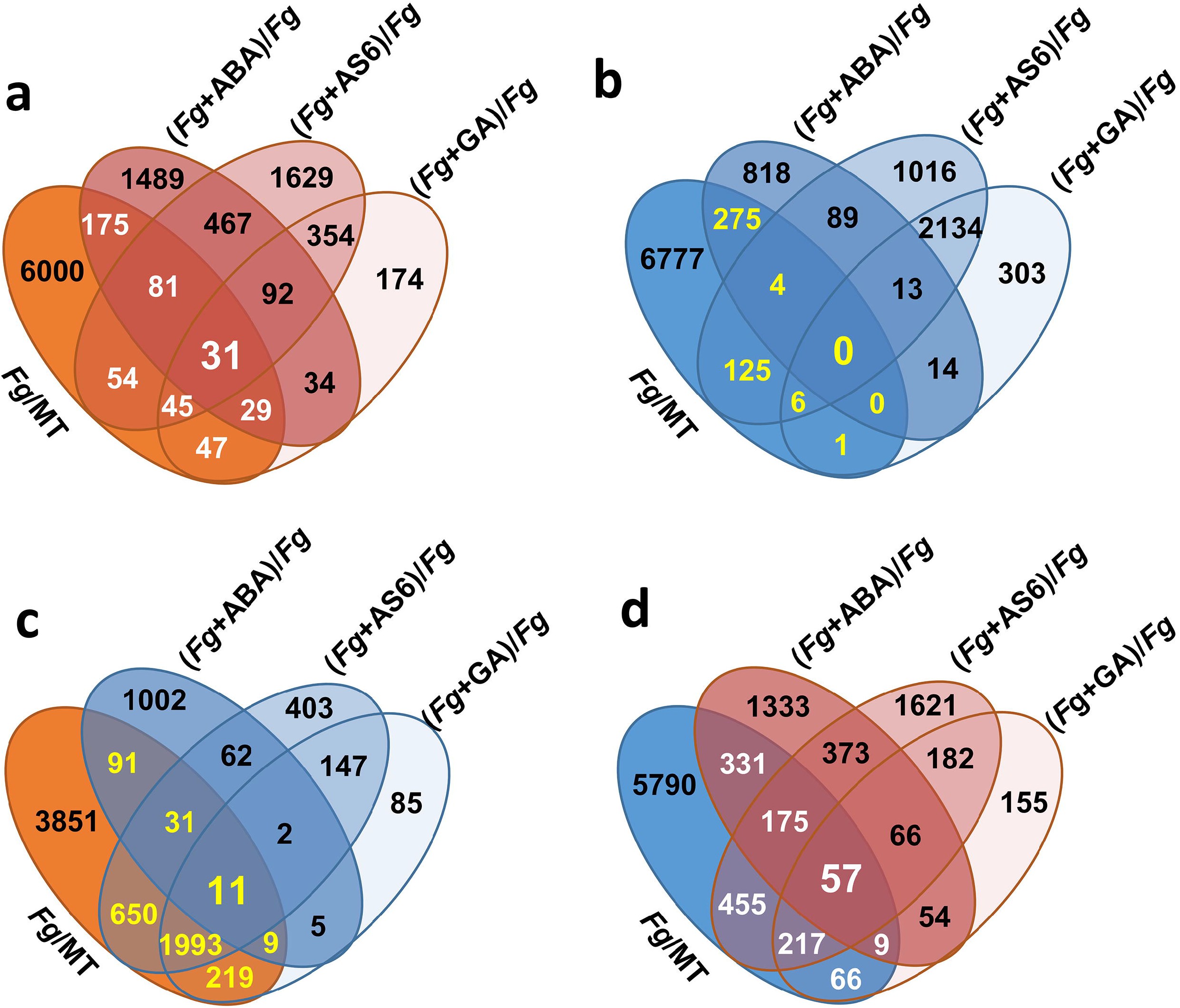
Share of *Fg* alone up- or down-regulated genes whose expression was enhanced or suppressed upon co-application of a phytohormone. In all instances, up-regulated genes are highlighted in shades of red, and down-regulated genes are highlighted in shades of blue. Enhanced genes are highlighted in white, suppressed genes are highlighted in yellow. The 6855 genes up-regulated by *Fg* (*Fg*/MT) whose expression was further enhanced **(a),** or suppressed **(c)** under the indicated co-applied conditions. Similarly, among the 7100 genes down-regulated by *Fg* that were further suppressed **(b),** or enhanced **(d),** under the indicated co-applied conditions. An annotated list of the 31 genes up-regulated under all conditions is available in **Additional File 8.**

At this global transcriptomic level, evidence of AS6 antagonism on wheat ABA receptors was not clearly observed; therefore, comparisons of DEGs **(Additional File 5: Figure S7)** and DEFE patterns **(Table 2**) elicited by co-applied *Fg*+AS6 and *Fg*+ABA, compared to *Fg* alone, were conducted. Of the 316 DEGs up-regulated by *Fg* alone and further enhanced by *Fg*+ABA, 11 DEGs were inhibited by *Fg*+AS6 (highlighted in **Additional File 5: Figures S6C).** These included a γ-glutamyl phosphate reductase (TraesCS3B01G395900) and a eukaryotic peptide chain release factor subunit 1-1 (TraesCS1A01G235000) found in the DEFE patterns M111100 and M111101 **(Additional File 5; Figure S8).** The expression of these two genes were highly significantly different between *Fg*+AS6 and *Fg*+ABA (adjusted p < 10^−11^, **Additional File 5: Figure S9).**

Conversely, among the 279 DEGs down-regulated by *Fg* and further repressed by *Fg*+ABA, 29 DEGs were enhanced by *Fg*+AS6 **(Additional File 5: Figure S6D).** These included four DEGs that were repressed by ABA in the presence or absence of *Fg* challenge (*Fg*+ABA or ABA treatments) as evidenced by DEFE pattern M202202 and M222202 **(Additional File 5: Figure S10).** The expression of these four genes were also highly significantly different between *Fg*+AS6 and *Fg*+ABA (adjusted p < 10^11^, **Additional File 5: Figure S9).** They include a BDX gene (TraesCS7B01G353600), orthologous to *Arabidopsis* AT4G32460; a transmembrane protein (TraesCS7A01G453100), orthologous to *Arabidopsis* AT5G11420; a leucine-rich repeat receptor-like protein kinase family protein (TraesCS3D01G040100), orthologous to *Arabidopsis* AT4G08850; and a nitrate transporter 1.1 (TraesCS1B01G225000). This result demonstrates that the putative ABA receptor antagonist AS6 exerted opposing effects on the wheat transcriptome as compared to ABA and highlights affected genes for future investigation.

### Wheat host phytohormone biosynthesis and signaling gene expression is altered by *Fg* challenge and phytohormone application

Targeted analysis of transcriptomic changes to putative ‘Fielder’ phytohormone biosynthetic and signaling pathways elicited by *Fg* challenge and phytohormone application were performed based on local sequence alignment of known homologs against the wheat DEGs. Wheat challenged with *Fg* alone elicited DEGs in classic phytohormone defense responses including up-regulating SA (PAL), JA (*OPR3*) and ET (ACS) biosynthetic genes and down-regulating a SA (*NPR1*) and repressive JA (JAZ) and up-regulating JA (*OCRA3*) and ET (*ETR1*) signaling genes **(Additional File 5: Figure S11, Additional File 1 Tabs ‘S2’ & ‘S3’).** *Fg*-challenge also impacted wheat biosynthetic and signaling pathways of non-defense phytohormones **(Figure 7)** with some mixed modulation of key contributors to the mevalonate and terpenoid biosynthesis pathways that feed into ABA, GA, CK and BR biosynthesis observed. Finally, *Fg* challenge down-regulated ABA biosynthetic genes (*β-CRTZ, AOX,* and *A8H*), while up-regulating two GA biosynthetic genes (*GA20ox* and *GA2ox*) and down-regulating a GA signaling repressor (*DELLA;* **Additional File 5: Figures S12 & S13).**

**Figure 7:**
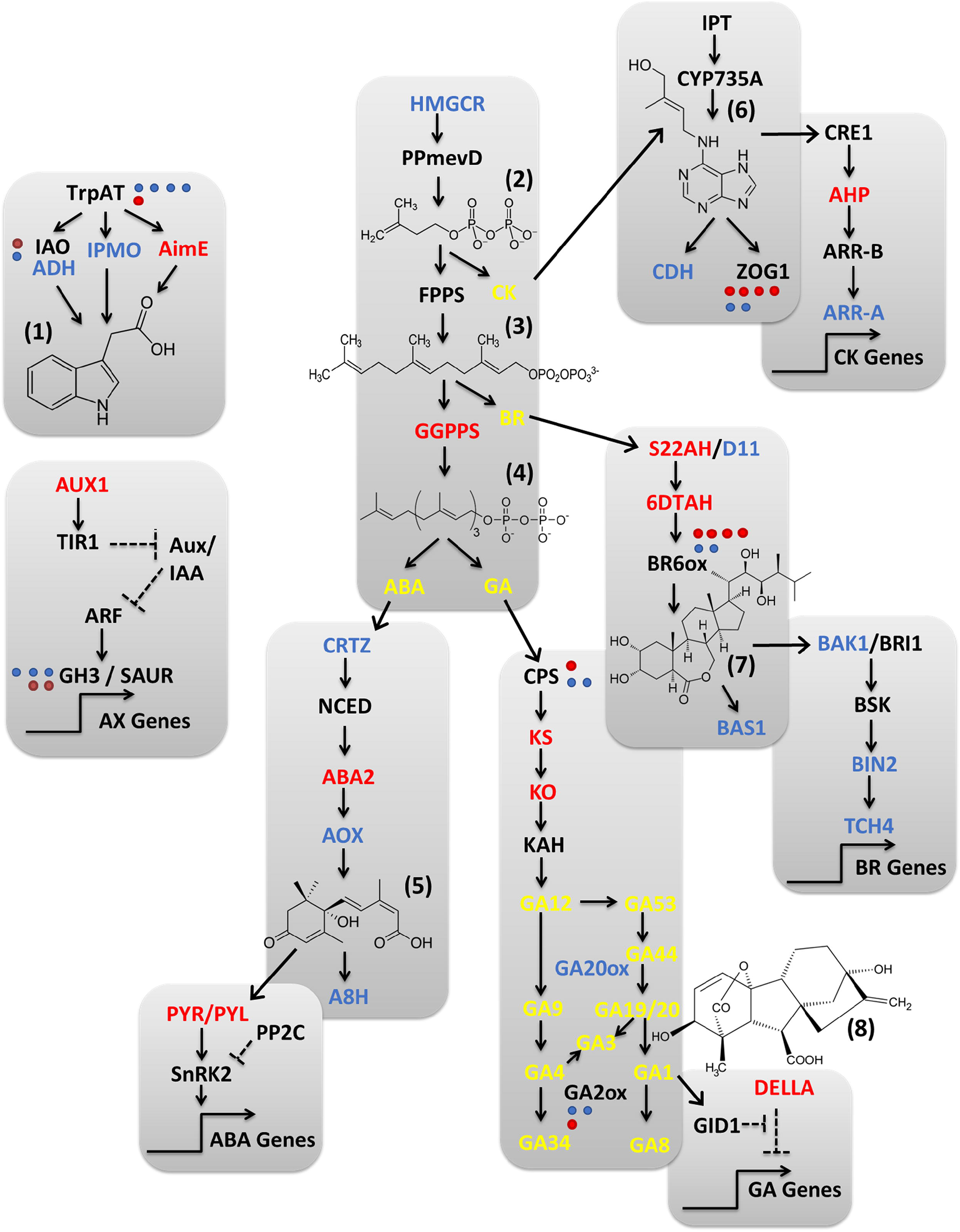
Effect of *Fg* infection on expression of other (non-classical defense) ‘Fielder’ phytohormone biosynthesis and signaling pathway genes. Chemical structures are shown for (1) indole acetic acid, (2) isopentenyl diphosphate (3) farnesyl diphosphate (4) geranylgeranyl diphosphate (5) abscisic acid (6) trans-zeatin (7) brassinolide and (8) giberellin (GA3). Compound acronyms are highlighted in yellow. ‘Fielder’ gene acronyms are represented as up-regulated (red; log_2_FC > 1.5) and down-regulated (blue; log_2_FC < −1.5), respectively (adj p < 0.01) compared to mock treatment. Annotations were based on Blast analysis selecting for transcripts with > 50 % sequence identity to known phytohormone signaling pathway members as annotated in the KEGG database. See **Additional File 1, Tabs S2 and S3** for detailed expression data.

The co-applied condition of *Fg*+ABA elicited an upregulation of wheat DEGs in the ABA (*HMGCR,* the rate-limiting *NCED, A8H*), GA (*GA2ox* and *GA20ox*), ET (*ACO*) and JA (*OPR3*) biosynthetic pathways **(Additional File 5: Figures S12 & S13).** This induction of GA20ox and GA2ox wheat genes was consistent with the observed change in GA metabolites when comparing *Fg* challenged and *Fg*+ABA treated wheat cultivar ‘Fielder’ phytohormone profiles **(Table 4).** The *Fg*+ABA condition also yielded significant modulation of ABA signaling genes, including both suppression of multiple receptors and enhanced expression of multiple negative regulator PP2Cs, and upregulation of SA (*PR1*), ET (*ETR1*), and BR (*TCH4*) signaling gene expression. Meanwhile, the co-applied *Fg*+GA suppressed the expression of GA biosynthetic genes (*KO* and *KAH*) and while also suppressing the negative regular *DELLA* involved in GA signaling. This condition also suppressed expression of ET (ETR1) and CK signaling genes (*A-ARR* and *AHP*) and BR biosynthetic genes (S22AH and *BR6ox*). The absence of an impact on wheat ABA signaling gene expression by the *Fg*+GA condition, and conversely GA signaling by the *Fg*+ABA condition, is generally consistent with the above noted suppression of *FUS3,* an important regulator at the intersection of ABA and GA crosstalk [32], under all conditions with *Fg* challenge **(Additional File 5: Figure S1).** Finally, the co-applied condition of *Fg*+AS6 up-regulated wheat DEGs in SA (*PAL*), JA (*OPR3*), ET (*ACS and ACO*), GA (*CPS, KS,* and *GA2ox*), and CK (*CDH* and *ZOG1*) biosynthesis while repressing genes in BR biosynthesis (*S22AH* and *BR6ox*). This condition also impacted wheat signaling gene expression with induction of SA (*NPR1*), JA (repressive *JAZ and ORCA3*), and BR (*TCH4*) while repressing ET (*ETR1*), GA (the repressive *DELLA*), and CK (*CDH* and *ZOG1*) genes alone **(Figure 8).** Interestingly, *Fg*+AS6 did not elicit changes to ABA metabolic or signaling pathways nor antagonistic changes to phytohormone pathways when compared to *Fg*+ABA (with the exception of DEGs corresponding to ACS in the ET pathway).

**Figure 8:**
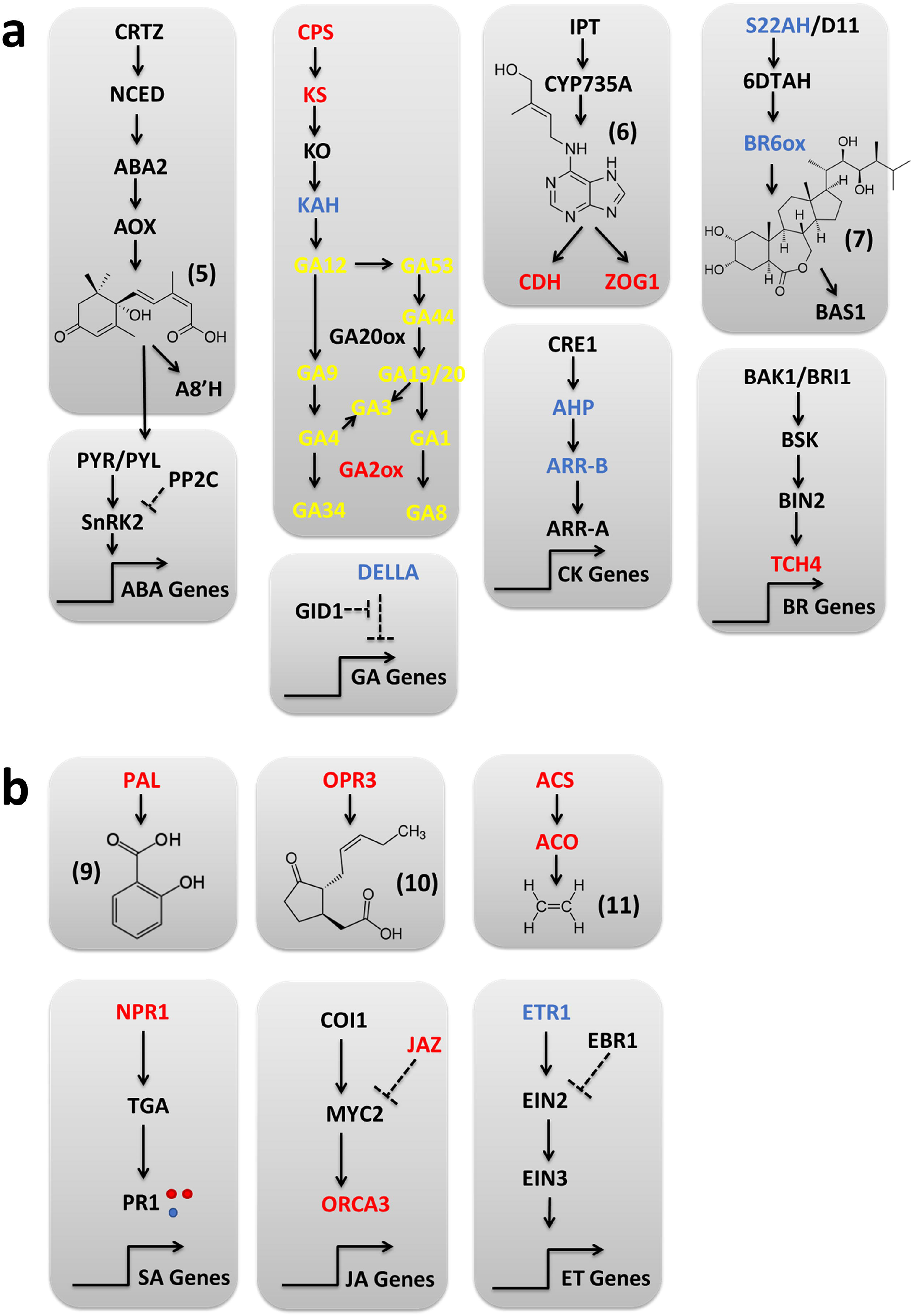
Phytohormone biosynthetic and signaling responses arising from co-application of AS6 at the time of *Fg* infection, with DEGs calculated compared to *Fg* alone. **a)** ABA and other affected (non-classical defense) hormone pathways with chemical structures for (5) abscisic acid, (6) trans-zeatin and (7) brassinolide. GA metabolite acronyms are highlighted in yellow. **b)** Affected classical defense pathways, with chemical structure for (9) salicylic acid (10) jasmonic acid and (11) ethylene. Legend details are otherwise the same as in Figure 7. See **Additional File 1, Tabs S2** and **S3** for details.

**Table 4.**
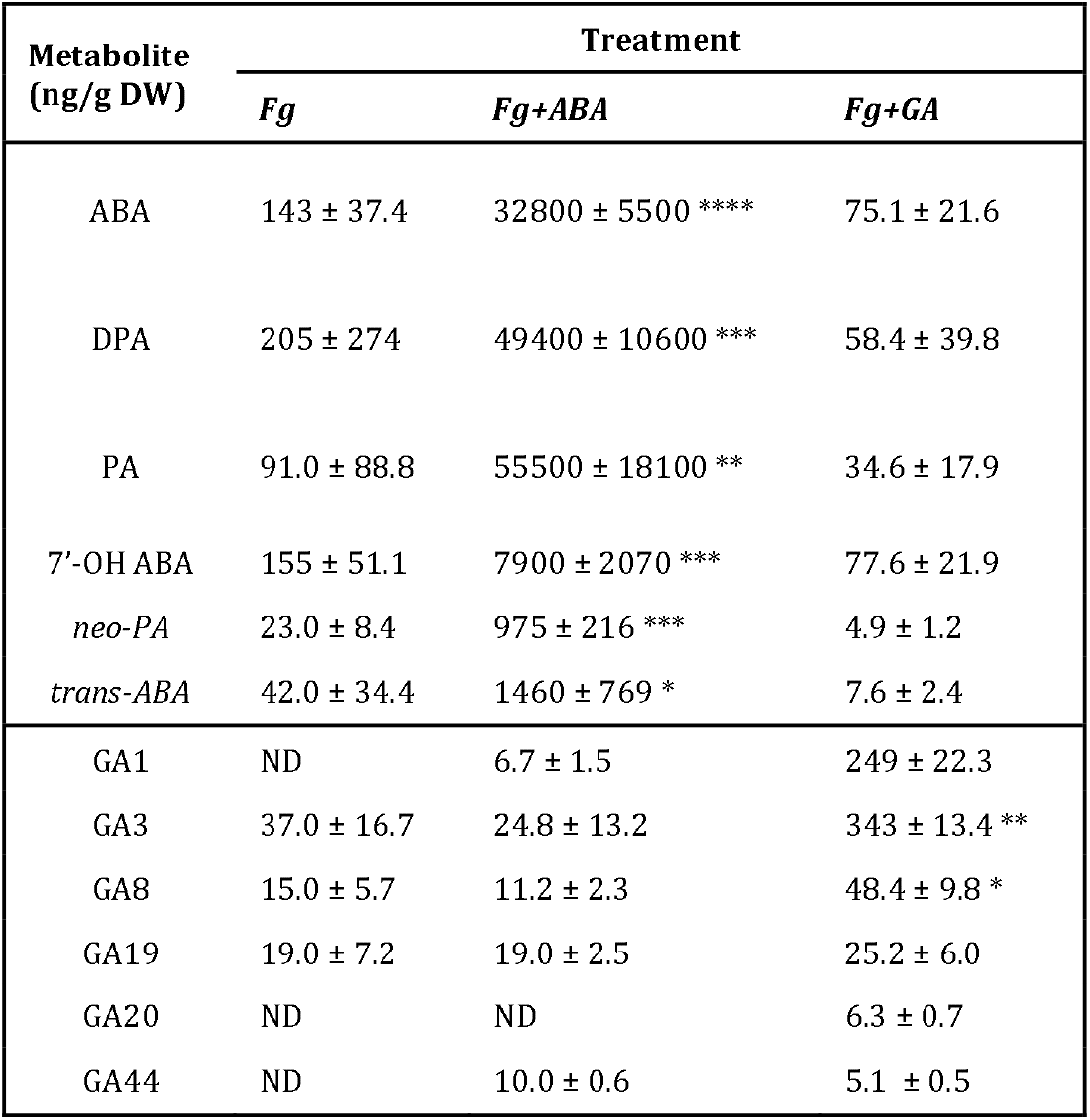
ABA and GA phytohormones and their metabolites detected in ‘Fielder’ spikes upon *Fg* challenge. Phytohormones and their associated metabolite levels were detected in ‘Fielder’ spikes (normalized to dry weight (DW)) inoculated with *Fg* in the absence (*Fg*) or presence of either ABA (*Fg* + ABA) or GA (*Fg* + GA). The ABA metabolites phaseic acid and dihydrophaseic acid are abbreviated PA and DPA, respectively, while undetected metabolites are denoted ‘ND’. Phytohormone changes were evaluated with one-way ANOVA with Dunnett post-hoc comparisons (* p ≤ 0.05, ** p ≤ 0.01, *** p ≤ 0.001, **** p ≤ 0.0001).

| Metabolite<br>(ng/g DW) | Treatment |  |  |
| --- | --- | --- | --- |
|  | *Fg* | *Fg*+ABA | *Fg*+GA |
| ABA | 143 ± 37.4 | 32800 ± 5500 **** | 75.1 ± 21.6 |
| DPA | 205 ± 274 | 49400 ± 10600 *** | 58.4 ± 39.8 |
| PA | 91.0 ± 88.8 | 55500 ± 18100 ** | 34.6 ± 17.9 |
| 7'-OH ABA | 155 ± 51.1 | 7900 ± 2070 *** | 77.6 ± 21.9 |
| *neo*-PA | 23.0 ± 8.4 | 975 ± 216 *** | 4.9 ± 1.2 |
| *trans*-ABA | 42.0 ± 34.4 | 1460 ± 769 * | 7.6 ± 2.4 |
| GA1 | ND | 6.7 ± 1.5 | 249 ± 22.3 |
| GA3 | 37.0 ± 16.7 | 24.8 ± 13.2 | 343 ± 13.4 ** |
| GA8 | 15.0 ± 5.7 | 11.2 ± 2.3 | 48.4 ± 9.8 * |
| GA19 | 19.0 ± 7.2 | 19.0 ± 2.5 | 25.2 ± 6.0 |
| GA20 | ND | ND | 6.3 ± 0.7 |
| GA44 | ND | 10.0 ± 0.6 | 5.1 ± 0.5 |

### Generating a consensus model of wheat transcriptomic responses to *Fg* challenge

In this work, the comparative transcriptomic responses of ‘Fielder’ are characterized upon phytohormone treatment in the presence or absence of *Fg* challenge. These transcriptome profiles were derived from high-quality RNA (RIN of 8.9 ± 0.38; **Additional File 1 Tab ‘S1’)** sequenced with a depth consistent with or greater than similar cereal-*Fusarium* reports (27 ± 7.7 million reads per sample), ultimately resulting in strong correlation between the five biological replicates investigated per treatment type **(Figure 1).** However, it remains unclear what proportion of, or trends in this high-quality ‘Fielder’ transcriptome data have the greatest biologic implications, especially considering that many transcriptomic investigations of cereal-*Fusarium* interactions do not agree between reports [6].

The transcriptomic results identified herein were compared with the four FHB susceptible genotypes ‘Roblin’ [10,11, 33], ‘Shaw’ [8], ‘Stettler’ and ‘Muchmore’ [9] in order to identify the strongest, consistent responses that may contribute to a consensus model of FHB infection. To draw this consensus, the wheat DEGs and trends highlighted here are discussed if two or more other genotypes also reported significant expression changes in spikelet tissue. First, of the approximately 2600 ‘Fielder’ key hub genes identified upon *Fg* challenge **(Additional File** 2 **Tab ‘WheatDEGs’@Col-P),** many were highly regulated by all genotypes. These changes include downregulation of fatty acid hydroxylase superfamily (13 DEGs with mixed regulation of 5 additional DEGs), photosystem II 22 kDa (3 DEGs), PISTILLATA-like MADS-box transcription factor (2 DEGs) and D-glycerate dehydrogenase/ hydroxypyruvate reductase DEGs and upregulation of C2 domain-containing protein-like (5 DEGS with mixed regulation of 4 additional DEGs) and yellow stripe-like transporter 12 (4 DEGs) **(Additional File 9 Tab ‘Hub-Genes’).** Among the genes involved in the phytohormone biosynthetic pathways **(Additional File 1, Tab S2),** *Fg* challenge up-regulated ET (three ACS synthase genes), JA (OPR3), and SA (PAL) biosynthetic DEGs while also up-regulating JA (ORCA3) and auxin (CH3) signaling DEGs across several genotypes **(Additional File 1, Tab S3).** Interestingly, pathogen treatment consistently repressed auxin biosynthesis (IPMO and ADH). There is also moderate agreement within the other comparative reports describing the upregulation of jasmonate ZIM-domain (7 DEGs) and downregulation of pectinesterase (4 DEGs), which are contradicted herein. Finally, an overlapping set of DEGs involved in aromatic and nitrogen metabolism are down-regulated and include a CTP synthase, DNA-directed RNA polymerase, and phenylalanine ammonium lyase along with mixed changes among several other DEGs **(Additional File 9 Tab ‘aromatic’ and ‘Cell N’).** Thirteen additional up-regulated phenylalanine ammonium-lyase DEGs involved in cinnamic acid biosynthesis demonstrate substantial agreement between genotypes **(Additional File 9 Tab ‘cinnamic’).**

In the remainder of wheat DEGs highlighted in this work, a consensus pattern emerged with GA treatment down-regulating and *Fg* (in the presence and absence of other treatments) up-regulating sets of DEGs. These DEGs were involved in heme (hemoglobin and cytochrome P450), oxidoreductase (cytochrome P450, polyphenyl oxidase, and aldehyde oxidase), and hydrolase (α/β hydrolase superfamily) activities **(Additional File 9 Tabs ‘Heme’, ‘oxidoreductase’, and ‘hydrolase’).** However, there is moderate agreement in comparative genotypes that are contradicted here, specifically with upregulation of hydrolase (β-amalase, β-fructofuranosidase, β-glucosidase, pectin acetylesterase, pectinesterase, and ribonuclease) and oxidoreductase (protochlorophyllide reductase, laccase, and respiratory burst oxidase) related DEGs **(Additional File 9 Tabs ‘hydrolase’ and ‘oxidoreductase’).**

## DISCUSSION

Comparative transcriptomic studies continue to be insightful in understanding general mechanisms modulating host resistance and susceptibility when challenged with *Fg* (reviewed in [6]). An encouraging flurry of recent publications emphasize roles for non-classical defense phytohormones in both resistance and susceptibility ([7–11], suggesting new focused alternative mechanisms underlying the host-pathogen interaction). To date, these studies have highlighted variable responses between examined cultivars, an aspect which is further complicated by recent evidence suggesting that non-classical defense phytohormones are also biosynthesized by *Fg* itself [19, 21, 23, 24]. In this light, the expression of a putative fungal ABA biosynthetic cytochrome P450 homolog under all conditions containing *Fg* examined herein is noted **(Additional File 5: Figure S2B),** lending further support to the role for fungal-derived ABA as an effector of infection, as has been proposed previously [19, 34].

### Assimilating ‘Fielder’ transcriptomic changes by comparing patterns of explicit features

The wheat whole-transcriptome analyses reported here were aimed at characterizing treatment-elicited changes in ‘Fielder’ and *Fg* gene expression by distilling the tens of thousands of observations into distinct strings of DEG permutations using DEFE. These combinations of explicit characteristics may then be used to understand the relationship between gene regulation and phenotype, especially when considering multiple treatment types [27]. DEFE descriptors quantifying the veritable impact of *Fg* challenge on wheat gene expression, highlighted abundant ‘Fielder’ expression changes elicited by ABA or GA application alone, and described the unexpected DEG overlaps of *Fg*+AS6 with *Fg*+GA (86 %) and *Fg*+ABA (23 %). Furthermore, by comparing individual DEFE descriptors from single phytohormone treatments with the intersection of several treatments (e.g., ABA⍰*Fg*⍰[*Fg*+ABA]), DEGs elicited by ABA or GA alone were shown to be common in the co-application samples in 91 % and 83 % of DEGs, respectively. By combining DEFE with topology overlap and gene association network analyses, more than 9500 ‘Fielder’ gene expression changes were characterized as *Fg*-related where the most highly enriched genes are putatively involved in stress response (alkaline shock protein 23, photosystem II 22 kDa, glyoxal oxidase, jasmonate ZIM domain proteins, and pectinesterase), cell structural integrity (fimbrin-like protein 2, glyoxal oxidase, and pectinesterase), membrane/lipid homeostasis (photosystem II 22 kDa and fatty acid hydroxylase), or molecular transport (clathrin assembly family protein, fimbrin-like protein 2, C2 domain-containing protein, and yellow stripe-like transporter). As expected in ‘Fielder’ spikelets, many of these enriched genes are putatively involved in flowering (PISTILLATA-like MADS-box transcription factor, fimbrin-like protein 2, pollen allergen, and pectinesterase) [35–37]. Finally, *FUS3* was identified as one notable gene in the network, having greater than 2000 connections. The *FUS3* ortholog in *Arabidopsis thaliana* (AT3G26790) is known to be involved in positive regulation of ABA biosynthesis and negative regulation of GA biosynthesis [32]. As *FUS3* gene expression was strongly suppressed across all *Fg*-challenged conditions, the presence of the pathogen likely dysregulates ABA and GA crosstalk and may decouple their often-oppositional biological phenotypes. In additional to biosynthesis, the disruption of both ABA and GA signaling has been observed in the presence of *Fusarium* virulence factors [38], suggesting there may be multiple mechanisms used by the pathogen to dysregulate ABA and GA related plant responses.

### Wheat transcriptomic responses to Fg challenge include ubiquitous traditional defense responses and more nuanced changes in non-classical defense phytohormones

*Fg*-challenge of wheat varieties elicits an incontestable shift of host gene expression with noted variations in the expression of phytohormone biosynthetic and signaling pathway genes [8, 9, 10, 11, 39]. When investigating ‘Fielder’ in this work, *Fg* challenge up-regulated both SA and JA biosynthetic pathways, notably with markers of late SA signaling highlighted, consistent with previous reports [8, 39]; while mixed differential gene expression changes of ET biosynthesis are juxtaposed to induction of one ET receptor gene, together suggesting ET pathways are being ‘primed’ for response. Beyond these classical defenses, *Fg*-challenge also elicited mixed expression changes of biosynthetic and signaling pathways of ABA, GA, IAA, BR, and CK where GA and IAA biosynthetic genes were up-regulated, GA and BA signaling genes were down-regulated, and ABA signaling genes were up-regulated.

When investigating cereal-*Fusarium* interactions using transcriptomic approaches, a variety of phytohormone responses have been observed which often do not agree between reports [6]. Wang et al. [9] may be most comparable to this work in wheat treatment, methodology (RNA-sequencing rather than microarray), and research focus (comprehensive phytohormone pathway regulation). Therefore, a direct comparison to Wang et al. [9] may be more appropriate than a field-wide discussion in which little consensus is noted. Both works characterize upregulation of genes involved in SA biosynthesis and SA, JA, ET, and ABA signaling. However, here we report increased JA and mixed ET biosynthetic gene expression changes, while Wang et al. characterize susceptible lines down-regulating JA and up-regulating ET biosynthetic genes with a contrasting increase in JA phytohormone concentration. Based on the consensus model of early biotrophic SA followed by later stage necrotrophic JA/ET responses [6, 8, 9], these differences may suggest that ‘Fielder’ tissues were analyzed while transitioning to the later JA (increased biosynthesis) / ET (‘primed’ biosynthesis) while Wang et al.’s ‘Settler’ and ‘Muchmore’ varieties were characterized at an earlier infection state. This hypothesis is consistent with the ‘Fielder’ transcriptome being derived from early-stage (24 hpt), challenged spikelets while the ‘Settler’ and ‘Muchmore’ transcriptomes were derived from three pooled whole spikes with singly challenged spikelets (4 days post treatment). It is also interesting that ‘Fielder’ may be more susceptible to FHB and thus not able to optimally utilize these distinct phases of phytohormone defense. In support to this hypothesis, ‘Settler’ and ‘Muchmore’ are red spring wheat commercial cultivars that have been characterized as “moderately susceptible” [40] while ‘Fielder’ is a white spring wheat that is used as a model system with equivalent susceptibility as the commercial cultivar ‘AC Crystal’ [41], which itself has been characterized to have “very poor” resistance to FHB [42]. Finally, the variations of phytohormone biosynthetic and signaling gene expression changes elicited by *Fg* may be, in part, due to the level of inspection. Herein, DEGs in these pathways are examined based on sequence similarity to orthologs with annotated function, while Wang et al. [9] and other reports have used GO term binning. By mapping individual genes, select genes could be monitored with greater precision, but global pathway changes may be at too high granular with many mixed results. Alternatively, by using GO terms alone, consensus or averaged gene expression changes in a pathway would be more clearly defined but more precise data for individual genes of interest would require additional investigation.

### ABA induces wheat DEGs observed in Fg infected tissues and may promote disease severity by misregulating phytohormone defense response and cell wall fortification mechanisms

Unlike the model of early SA and later JA/ET responses triggered in wheat challenged with *Fg* [6, 8, 9], the gene expression changes upon application of individual phytohormones to wheat has not been coalesced into a single model. Perhaps the most comprehensive study of the gene expression changes has been the work of Qi et al. [11] where wheat spikelets were treated with IAA, GA, ABA, ET, CK, SA or JA (methyl jasmonate) and resultant gene expression changes were characterized by microarray. Surprisingly, there were many contradictory findings when comparing the findings of Qi et al. and the transcriptomic changes presented here, namely in the number of DEGs identified per treatment and the relative proportionality between up- and down-regulated DEGs within a given treatment. These differences may be in part due to comparison between pooled wheat spikes vs treated spikelets or the comparison between two different genotypes, ‘Roblin’ vs ‘Fielder’. It may also be method-dependent where wheat microarray analysis is limited to 50,000 – 60,000 genes [11,39]; and additional stringency criteria must be applied to properly control for complications such as cross-hybridization and limited dynamic assay range (as reviewed in [43]). This hypothesis is supported by wheat microarray studies which typically report 100 – 3500 DEGs [11, 39, 44, 45], while wheat RNA-seq studies typically report 25,000 – 40,000 DEGs (this work; [8, 9]).

Despite the tissue and methodological differences between this work and Qi et al. [11], both studies report a significant overlap between gene expression changes elicited by *Fg* challenge and ABA application as well as the ABA-dependent regulation of polyphenolic compound-related gene expression. Herein, exogenously applied ABA alone yielded a general repression in ‘Fielder’ where 57% of up-regulated and 75% of down-regulated genes were also elicited by *Fg* challenge. The vast majority of the genes (> 90 %) significantly modified by ABA alone and *Fg* alone were also significantly modified in the co-applied *Fg*+ABA condition, highlighting the relevance of findings from studies of ABA-alone with respect to understanding the interaction of ABA with *Fg* infection. However, ABA treatment outcomes are not entirely overlapping with those elicited by either *Fg* or *Fg*+ABA. One such notable case is the ABA down-regulated but *Fg* up-regulated polyphenolic compound related genes. Phenylpropanoid metabolism is involved in biosynthesis of lignins, cinnamic acids, and flavonoids that are in turn used for cell wall fortification and *Fg* defense [46–50]. FHB resistant wheat and durum varieties have been shown to have key anatomical differences, improved cell wall structure including the additional lignin deposition, and greater accumulation of the cinnamic acid metabolite p-coumaric acid in infected tissues as compared to susceptible varieties [47, 51–53]. Furthermore, some polyphenolic compounds have documented anti-fungal activities [40, 54]. In addition to the ABA alone treatment, the co-applied *Fg*+ABA treatment also highlighted altered physical defense responses and non-classical defense phytohormone signaling specifically suppressing IAA and enhancing BR signaling. IAA signaling suppression is consistent with heightened classical defense responses and has been detected in susceptible near isogenic lines of the 2DL QTL, compared to more resistant lines [55, 56]. Enhanced BR signaling included the expression of a cell wall re-modelling xyloglucan endotransferase gene (*TCH4*) that may contribute to FHB susceptibility through inappropriately regulated cell wall modification. Together, wheat transcriptomic changes elicited within ABA treatment groups suggests this phytohormone promotes FHB disease severity [19, 20] by inducing wheat gene expression changes necessary for *Fg* infection including misregulating classical defense phytohormone signaling and physical defense mechanisms such as plant cell wall fortification and polyphenolic metabolism.

### GA induces global metabolic shifts in both the wheat host and Fg pathogen

Historically, both genetic and chemical approaches have been applied to modulate GA metabolism for the improvement of agronomic wheat characteristics [57, 58]. It remains to be determined whether targeting GA metabolism could be a reasonable strategy for controlling FHB based on the observation that GA-regulated wheat anthesis and plant height are strongly correlative with infection and disease severity [1, 59–61]. Based on the transcriptomic profiles of wheat treated with diverse phytohormones, it was determined that ABA and GA elicit the strongest transcriptional antagonism of any phytohormone pair, with approximately 40% of their DEG being antagonistic [11]. These findings are too in agreement with previous reports of ABA promoting and GA suppressing FHB disease severity [19, 20]. Here, exogenously applied GA alone resulted in a relatively even mix of up-regulated and down-regulated ‘Fielder’ genes including nearly 500 gene expression changes that directly opposed those elicited by *Fg* alone. GO enrichment suggest that these oppositional effects included down-regulated secondary metabolism and defense response genes, coincident with up-regulated primary metabolic processes. Analysis of *Fg*+GA subsets notably highlighted suppression of genes involved in flower development. These GA derived DEGs describe a broader metabolic switch with moderate overlap compared to the primary carbohydrate, TCA, and nitrogen metabolic changes elicited by DON treatment [62]. It is also interesting to note that *Fg* transcriptionally regulates its own metabolism when transitioning from a biotrophic to necrotrophic state during the first 48 hours post infection [63]. In Buhrow et al., [20], the effect of *Fg*+GA co-application on *Fusarium* gene expression was characterized with this phytohormone eliciting DEG broadly within amino acid, carbohydrate, and lipid metabolism and specifically regulating five genes in inorganic nitrogen or amino acid nitrogen metabolism. This coupling of GA and nitrogen metabolism has been previously described in the related *Fusarium moniliforme* [64]. Therefore, these findings suggest that GA may limit FHB severity through global metabolic changes on both the wheat host and *Fg* pathogen rather than through a select few defense pathways.

### The *Fg*+AS6 treatment functions independently of ABA-signaling to antagonize ‘Fielder’ gene expression elicited by ABA⍰*Fg*⍰[*Fg*+ABA] treatments

A rationally designed ABA analog with well-characterized ability to disrupt ABA receptor interactions in dicots (AS6; [25]) was previously reported to elicit no observable FHB phenotypic effect when co-applied with *Fg* challenge [20]. It remains unclear whether AS6 reaches its intended target (due to modification or degradation), is potent against wheat ABA-receptor interactions, or elicits a response which is sustained as FHB phenotype is monitored. However, comparative transcriptomics was applied to clarify whether AS6 (*Fg*+AS6) elicits responses antagonistic with those of ABA (*Fg*+ABA). *Fg*+AS6 treatment both up- and down-regulated *F. graminearum* pathogen gene expression with more than 550 antagonist effects compared to *Fg*+ABA. *Fg* has been proposed to biosynthesize ABA *de novo* ([19, 34]) but lacks clearly identifiable ABA receptors through which ABA-specific antagonism would be triggered. When focusing on the ‘Fielder’ transcriptome, *Fg*+AS6 had no effect on ABA biosynthetic or signaling gene expression, arguably the metabolic pathway expected to be most affected by a potent ABA receptor antagonist. This lack of antagonism may be partially attributed to differences in specificity between ‘Fielder’ and dicot ABA receptors. In fact, ABA receptors in the wheat cultivar ‘Thatcher’ have been reported to have unique selectivity compared to *Arabidopsis* receptors across a panel of ABA analogs [65]. However, *Fg*+AS6 did elicit gene expression changes to ‘Fielder’, sometimes in agreement with *Fg*+ABA and other times in agreement *Fg*+GA treatment, including up-regulating genes in the SA, JA, and ET defense response pathways.

Most notably, *Fg*+AS6 elicited robust antagonist responses compared to ABA⍰*Fg*⍰[*Fg*+ABA]. This treatment mitigated the upregulation ofγ-glutamyl phosphate reductase and eukaryotic peptide chain release factor subunit 1-1 genes elicited by ABA⍰*Fg*⍰[*Fg*+ABA]. The expression of γ-glutamyl phosphate reductase has been tied to abiotic stress response [66]; while eukaryotic peptide chain release factor subunit 1-1 has been tied to plant growth and development including appropriate cell-wall lignification [67]. *Fg*+AS6 also robustly up-regulated gene expression of a leucine-rich repeat receptor-like protein kinase (LRRKs) and nitrate transporter 1.1 that were down-regulated by ABA⍰*Fg*⍰[*Fg*+ABA] treatments. There have been more than 500 wheat LRRKs annotated with function in innate immunity responses, SA signaling repression, and grain development with some members even being responsive to FHB and the presence of mycotoxins [68, 69]. LRRKs have also been linked to FHB resistance in durum [59]. Nitrate transporter 1.1 has been shown to transport nitrate and nitrite, peptides and amino acids, phytohormones (auxin, ABA, JA-Isoleucine, and GA) and glucosinolates (reviewed in [70]). This transporter has also been suggested to function as a bridge between phytohormone responses during pathogen stress (reviewed in [70, 71]). As all of the robust *Fg*+AS6 responses can be tied to plant stress responses, and in some cases even FHB and phytohormone pathways, AS6 may in fact promote FHB resistance in a manner that is ABA-signaling independent, and not observable upon a single early-stage treatment in the late-stage phenotype.

### Agreement between transcriptomic responses derived from several genotypes support a consensus model of the wheat-Fg interaction

Previous transcriptomics reports of *Fg*-challenged wheat highlighted up-regulated basal defenses, antagonism of pathogen-mediated modulation of phytohormone pathways, and classical phytohormone defense signaling [12–16] with an early biotrophic SA followed by later stage necrotrophic JA/ET responses [6, 8, 9]. However, many transcriptomic investigations of cereal-*Fg* interactions do not agree between reports [6]. To additively contribute to this consensus model, gene expression changes were compared between five FHB susceptible wheat genotypes. Generally ubiquitous upregulation of C2 domain-containing protein-like, jasmonate ZIM-domain, yellow stripe-like transporter 12, phenylalanine ammonium-lyases (cinnamic-acid metabolism) and downregulation of hydrolases that putatively cleave fatty acids, pectins, ribose, and sugar moiety substrates were observed. Interestingly, ABA treatment also regulates DEGs involved in cell wall-modifying polyphenolic and cinnamic acid metabolism and a saccharide hydrolase/ transferase, xyloglucan endotransferase. Meanwhile, GA treatment responses are targeted to metabolic processes like those elicited by DON treatment (discussed previously).

## CONCLUSIONS

This work explored the comparative transcriptomics of wheat ‘Fielder’ spikes treated with ABA, the ABA antagonist AS6, or GA in the presence and absence of *Fg* challenge. The findings reported herein emphasize the vast impact of pathogen challenge, even as early as 24 hpt, and highlight wheat genes broadly involved in defense responses, cell structural integrity, molecular transport, and membrane/lipid metabolism as strongly correlative with *Fg* pathogen challenge. Although ABA and GA often elicit oppositional plant phenotypes, the presence of *Fg* down-regulated *FUS3* for the crosstalk of these two phytohormones, supporting independent mechanisms by which ABA promotes and GA reduces FHB disease severity. ABA may promote disease by eliciting gene expression responses common to those elicited by the pathogen, misregulating defense responses by further exacerbating gene expression, and altering cell wall fortification including through polyphenolic metabolism. AS6 does not traditionally antagonize ABA signaling or biosynthesis in ‘Fielder’ but opposes ABA- and *Fg*-elicited responses including those in phytohormone defense signaling and transport and cell wall organization. GA application elicits a diverse metabolic reprogramming of both the wheat host and *Fg* pathogen during the early infection when *Fg* is transitioning from a biotrophic to necrotrophic lifestyle. This reprogramming may result in a less susceptible host and/or less virulent pathogen ultimately reducing FHB disease severity observed with GA co-application. Finally, the biological implications of these transcriptome responses on the wheat-*Fg* interaction were evaluated by assessing common findings across five wheat genotypes, ultimately additively contributing to a limited consensus model of disease response and severity.

## METHODS

### Chemicals, phytohormones and *Fg* inoculum preparation

Gibberellin A3 was purchased from Sigma-Aldrich (St. Louis, MO). The National Research Council Hormone Profiling Facility provided (+)-ABA, while 3⍰-hexasulfanyl-(+)-ABA was synthesized as described in Takeuchi et al., [25] and provided by Kenneth Nelson and Suzanne Abrams at the University of Saskatchewan. Phytohormone stocks were solubilized in deionized water as sodium salts by 1.0 N NaOH titration and stored at −20 °C in amber vials. Working solutions were made in deionized water and pH was adjusted to 7.0 ± 0.05. *Fg* GZ3639 [72] was propagated on potato dextrose agar (PDA; Sigma-Aldrich) at 25 ° C for five days. To obtain spores, carboxymethylcellulose (CMC) liquid media (1.5 % CMC (Sigma), 0.1 % NH4NO_3_, 0.1 % KH_2_PO_4_, 0.05 % MgSO_4_·7H_2_O, and 0.1 % yeast extract;) was inoculated with a marginal 5 mm square PDA plug and grown for five days at 27 ° C, shaking at 170 rpm. Macroconidia were isolated by filtering through one layer of cheese cloth and 25 μm Miracloth filter (EMD Millipore; Billerica, MA), washed three times with sterile water, and quantified using a haemocytometer and light microscopy.

### Propagation of plants, pathogen-challenge +/− co-applied phytohormones and phytohormone profiling

*T. aestivum* ‘Fielder’ was grown in Sunshine^R^ Mix 8 (Sungrow Horticulture, Agawam, MA) and maintained in climate-controlled chambers with a 16 h photoperiod, at 25 °C followed by 8 h of dark at 16 °C every day. Plants were watered as needed and fertilized biweekly with 20-20-20 (N-P-K). At the two-leaf stage plants were treated Intercept™ (Bayer Crop Science, Calgary, AB) as an aphid preventative, as previously described [20, 39]. During anthesis, all florets from a central spikelet were point inoculated with 10 μL of 5.0 x 10^4^ *Fg* GZ3639 macroconidial suspension or deionized water (mock). For phytohormone co-application, this inoculum was supplement with and without 1.0 mM ABA, GA3 or AS6. To promote infection, wheat plants were transferred to climate-controlled chambers with misting to achieve 90 % humidity for 24 h. Five entire spikes were then harvested from five different plants (five biological replicates) for each treatment group, each as a single tissue sample, and analyzed for phytohormone content. Collectively, there were 7 treatments (*Fg,* MT, ABA, GA, *Fg*+AGA, *Fg*+GA. *Fg*+AS6) and 35 samples in total. Individual spikes were flash frozen and ground in liquid nitrogen. Phytohormones were extracted from individual replicate spikes and quantified by UPLC/ESI-MS/MS at the National Research Council of Canada in Saskatoon, Canada, as described [73–77]. Phytohormone content differences in *Fg*-challenged spikes upon phytohormone application were analyzed with one-way ANOVA with Dunnett post-hoc comparisons using GraphPad Prism 6 (GraphPad Software, Inc. La Jolla, CA).

### RNA sequencing, data processing and differential expression analyses

Total RNA was extracted from the 35 samples above, purified, quantified and sequenced as described previously [20]. In short, RNA was purified using the RNeasy Plant Mini Kit (Qiagen, Mississauga, ON) and treated with DNaseI (Qiagen). RNA quality was evaluated using NanoDrop ND-8000 (NanoDrop, Wilmington, DE) and agarose gel electrophoresis. RNA library construction was completed using 1.0 μg total RNA and the TruSeq RNA Sample Prep Kit v2 (Illumina, SanDiego, CA). Library quality was assessed on the 2100 Bioanalyzer (Agilent Technologies Inc. Santa Clara, CA) equipped with a DNA 1000 chip. Library concentrations were determined by quantitative polymerase chain reaction (qPCR) using the KAPA SYBR FAST ABI Prism qPCR Kit (Kapa Biosystems, Wilmington, MA) and the StepOnePlus Real-Time PCR System (Applied Biosystems, Foster City, CA). RNA samples were multiplexed at a sequencing depth of five libraries per lane. Equimolar concentrations of the libraries were pooled and a final concentration of 12 pM was used for clustering in cBOT (Illumina) flowcell lanes. The samples were then sequenced (2×101 cycles, paried end reads) on the HiSeq2500 (Illumina) using the TruSeq SBS Kit v3-HS 200 cycles Kit (Illumina). The raw RNA-seq reads (available at GEO GSE137895) were preprocessed by trimming the adaptor sequences, filtering low-quality reads (Phred Score ≤ 20 [78]) and eliminating short reads (length ≤ 20 bps) using software package FASTX-toolkit [79]. After filtering, barcode and adaptor removal, an average of 27 million paired reads per sample was retained for subsequent read mapping through the RNA-seq data processing procedures. The IWGSC RefSeq v1.0 complete reference genome and corresponding annotation v1.0 were used as reference for the analysis of wheat RNA-seq data [26]. Following the recommendation of IWGSC, the chromosome partitioned version (161010_Chinese_Spring_v1.0_pseudomolecules_parts.fasta) was used and the gff3 file was reformatted accordingly. Only the high confidence gene models were considered in the mapping process. The *Fg* reference genome (*Fusarium graminearum* str. PH-1) was obtained from EnsemblFungi (http://fungi.ensembl.org/ Release 35). Wheat and *Fg* genomes and annotation data from both species were combined into a single reference [80]. This combined reference genome contains 124935 gene models, 14145 from *Fg* and 110790 from wheat. The cleaned RNA-seq reads in each sample were mapped using STAR v2.5.3a [81] to generate gene-level counts. DESeq2 [82] was used for data normalization and subsequent differentially expressed gene (DEG) analysis for each pairwise comparison. Normalized read counts along with log2 fold change and *p*-value, and adjusted *p*-values based on Benjamini and Hochberg procedure [83], were provided for downstream data analysis.

### Data reduction and partitioning

The output data from DESeq2 [82] was reduced in size to a set of differentially expressed genes. We applied the criteria of absolute log2FC ≥ 1, adjusted p ≤ 0.01 and one of the pair of compared samples has to be significantly expressed, i.e ≥ 10 reads. Six informative pairwise comparisons between each treatment and MT and three pairs between co-application of *Fg* with a phytohormone and *Fg* alone and a comparison between co-applications of *Fg* with ABA and with AS6 **(Figure 1d, Table 1**) were performed. The differential expression feature extraction method (DEFE; [27]) was applied to partition the DEGs into various feature patterns. We compiled the six treatments as compared with MT as one series of “M” patterns: 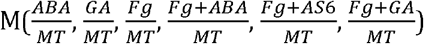. For example in the pattern W201110, at each comparison, “0” denotes no change, “1” for up- and “2” for down-regulation; this example pattern means a gene is down-regulated by ABA alone, no significant change with regard to GA, whether applied with or without *Fg*, but up-regulated by *Fg* alone, *Fg*+ABA and *Fg*+AS6. Similarly, we compiled the three combinations of respective phytohormone with *Fg* infection as compared with *Fg* infection alone in a series “F” patterns: 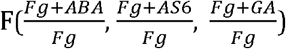.

### Clustering, correlation, network, gene ontology enrichment, and orthology analyses

These analyses were done following the same methods described in [8].

### Phytohormone pathway gene identification and analyses

BLAST analyses of representative biosynthetic and signaling query sequences were performed against the DEGs derived in this study based on the Kyoto Encyclopedia of Genes and Genomes (KEGG) Database (cut off > 50 % amino acid sequence identity). This yielded significant DEGs (log_2_FC > 1.5 or < −1.5; adj p < 10^−2^; to highlight the more highly modified) covering the spectrum of plant phytohormone biosynthesis and signaling pathways **(Additional file 1 Tabs ‘S2’ & ‘S3’** respectively).

## List of abbreviations

ABA: abscisic acid
BR: brassinosteroid
CK: cytokinin
DEFE: differential expression feature extraction
DEG: differential gene expression
*Fg*: *Fusarium graminearum*
FHB: Fusarium head blight
ET: ethylene
GA: gibberellic acid
IAA: indole acetic acid (auxin)
JA: jasmonic acid
SA: salicylic acid

## DECLARATIONS

### Ethics approval and consent to participate

Not Applicable

### Consent for Publication

Not Applicable

### Availability of data and materials

The datasets generated and/or analyzed during the current study are available in the GEO repository, GSE137895.

### Competing interests

The authors declare that they have no competing interests.

### Funding

This research was funded to M.C.L and Y.P. by the Canadian Wheat Improvement Program, National Research Council of Canada (grant # 008237, #011649)

### Author Contributions

LMB performed all experiments, as well as contributed to the design of experiments, data analysis, interpretation and writing of the manuscript. ZL and DC contributed to data processing and analysis. TS contributed to analysis of the fungal data in particular. NAF contributed to methods development, data analysis and writing of the manuscript. YP contributed to data processing, analysis, interpretation and writing of the paper. MCL conceived of the idea and contributed to experimental design, data analysis and interpretation and writing of the manuscript. All authors read and approved the final manuscript.

## Acknowledgements

We are grateful to Dr. L. Irina Zaharia at the Hormone Profiling Service, National Research Council of Canada, Saskatoon, Canada, for data collection and methodological input. We thank Dr. Susan McCormick at the United State Department of Agriculture for providing the *F. graminearum* strain GZ3639, Kenneth Nelson and Suzanne Abrams at the University of Saskatchewan for providing AS6. This manuscript represents NRC Communication # 58262.

## ADDITIONAL FILES

**Additional File 1** (xlsx): supplementary tables. This file contains three tables: (S1) RNA-seq read mapping statistics, (S2) ‘Fielder’ phytohormone biosynthetic pathways are affected by co-application of hormones with *Fg* challenge, compared to hormone- or *Fg-* challenge alone, and (3) ‘Fielder’ phytohormone signalling pathways are affected by co-application of hormones with *Fg* challenge, compared to hormone- or *Fg*-challenge alone.

**Additional File 2** (xlsx): Differentially Expressed Wheat genes. This file contains three worksheets: (1) a list of 26001 wheat DEGs, their expression values, cluster membership, gene network property, differential expression (log2FC and adj p values), DEFE pattern, and correlation analysis, etc.; (2) DEFE statistics, number of genes belong to each DEFE pattern, full lists of Tables 2 and 3; (3) correlation analysis of the 58 clusters.

**Additional File 3** (xlsx): Differentially Expressed Genes of *Fuzarium graminearum.* This file contains two worksheets: (1) a list of 4872 *Fg* DEGs, including their expression values, differential expressions (log2FC and adj p values), and DEFE pattern.

**Additional File 4** (xlsx): Enrichment index of various genes with their membership in various groups including DEGs, FHB-related genes, top 1% gene Network, Key hub genes.

**Additional File 5** (pdf): 13 Supplementary Figures.

**Additional File 6** (xlsx): Three series of Gene Ontology analyses in 32 worksheets plus a summary note worksheet: (1) FD series: ten lists of wheat genes down-regulated by *Fg*; (2) FU series: 19 lists of wheat genes up-regulated by *Fg*; (3) F series: three groups of *Fg* genes differentially expressed as affected by AS6 and others.

**Additional File 7** (xlsx): twenty-six up-regulated genes by *Fg* alone were suppressed by *Fg*+AS6

**Additional File 8** (xlsx): Thirty-one wheat genes up-regulated by *Fg* and further enhanced by co-application with all three hormones, respectively.

**Additional File 9** (xlsx): Comparisons across five wheat genotypes susceptible to FHB. This file contains seven worksheets: (1) network hub genes; (2) genes involved in the aromatic metabolism; (3) genes involved in the cellular nitrogen metabolism; (4) genes involved in the cinnamic acid biosynthesis; (5) genes involved in the heme activity; (6) genes involved in the oxidoreductase activity; and (7) genes involved in the hydrolase activity.

## Notes

https://www.ncbi.nlm.nih.gov/geo/query/acc.cgi?acc=GSE137895

### Competing Interest Statement

The authors have declared no competing interest.

## REFERENCES

1. Bai G, Shaner G. Management and resistance in wheat and barley to Fusarium head blight. Annu Rev Phytopathol. 2004;42:135–61.

2. Parry MAJ, Reynolds M, Salvucci ME, Raines C, Andralojc PJ, Zhu X-G, et al. Raising yield potential of wheat. II. increasing photosynthetic capacity and efficiency. J Exp Bot. 2011;62:453–467.

3. Foroud NA, Pordel R, Goyal RK, Ryabova D, Chatterton S, Kovalchuk, I. Chemical activation of the ethylene signalling pathway promotes wheat resistance to *Fusarium graminearum*. Phytopathology. 2019; 109:796–803.

4. Buerstmayr H, Ban T, Anderson JA. QTL mapping and marker-assisted selection for fusarium head blight resistance in wheat: a review. Plant Breed. 2009;128:1–26.

5. Cuperlovic-Culf M, Loewen M, Rajagopalan N, Surendra A. Perspectives on the specific targeting of *Fusarium graminearum* for the development of alternative head blight treatment approaches. Plant Pathol. 2017;66:1391–1403.

6. Kazan K, Gardiner DM. Transcriptomics of cereal-*Fusarium graminearum* interactions: what we have learned so far. Mol Plant Pathol. 2018;19:764–778.

7. Biselli C, Bagnaresi P, Faccioli P, Hu X, Balcerzak M, Mattera MG, et al. Comparative transcriptome profiles of near-isogenic hexaploid wheat lines differing for effective alleles at the 2DL FHB resistance QTL. Front Plant Sci. 2018;9:37.

8. Pan Y, Liu Z, Rocheleau, H, Fauteux F, Wang, Y, McCartney C, et al. Transcriptome dynamics associated with resistance and susceptibility against Fusarium head blight in four wheat genotypes. BMC Genomics. 2018;19:642.

9. Wang L, Li Q, Liu Z, Surendra A, Pan Y, Li Y, et al. Integrated transcriptome and phytohormone profiling highlight the role of multiple phytohormone pathways in wheat resistance against fusarium head blight. PLoS One. 2018;13:e0207036.

10. Brauer EK, Rocheleau H, Balcerzak M, Pan, Y, Fauteux F, Liu Z, et al. Transcriptional and hormonal profiling of *Fusarium graminearum*-infected wheat reveals an association between auxin and susceptibility. Physiol Mol Plant P. 2019;108: 33–39.

11. Qi PF, Jiang YF, Guo ZR, Chen Q, Ouellet T, Zong LJ, et al. Transcriptional reference map of phytohormone responses in wheat spikes. BMC Genomics 2019;20:390.

12. Rawat N, Pumphrey MO, Liu S, Zhang X, Tiwari VK, Ando K, et al. Wheat Fhb1 encodes a chimeric lectin with agglutinin domains and a pore-forming toxin-like domain conferring resistance to Fusarium head blight. Nat Genet. 2016;48, 1576–1580.

13. Kage U, Yogendra KN, Kushalappa AC. TaWRKY70 transcription factor in wheat QTL-2DL regulates downstream metabolite biosynthetic genes to resist *Fusarium graminearum* infection spread within spike. Sci Rep. 2017;7:42596.

14. Li G, Zhou J, Jia H, Gao Z, Fan M, Luo Y, et al. Mutation of a histidine-rich calcium-binding-protein gene in wheat confers resistance to Fusarium head blight. Nat Genet. 2019;51:1106–1112.

15. Su Z, Bernardo A, Tian B, Chen H, Wang S, Ma H, et al. A deletion mutation in TaHRC confers Fhb1 resistance to Fusarium head blight in wheat. Nat Genet. 2019;51:1099–1105.

16. Fauteux F, Wang Y, Rocheleau H, Liu Z, Pan Y, Fedak G, et al. Characterization of QTL and eQTL controlling early *Fusarium graminearum* infection and deoxynivalenol levels in a Wuhan 1 x Nyubai doubled haploid wheat population. BMC Plant Biol. 2019;19:536.

17. Chen X, Steed A, Travella S, Keller B, Nicholson P. *Fusarium graminearum* exploits ethylene signalling to colonize dicotyledonous and monocotyledonous plants. New Phytol. 2009;182:975–983.

18. Sun Y, Xiao J, Jia X, Ke P, He L, Cao A, et al. The role of wheat jasmonic acid and ethylene pathways in response to *Fusarium graminearum* infection. Plant Growth Regul. 2016;80:69–77.

19. Qi PF, Balcerzak M, Rocheleau H, Leung W, Wei Y-M, Zheng Y-L, et al. Jasmonic acid and abscisic acid play important roles in host-pathogen interaction between *Fusarium graminearum* and wheat during the early stages of Fusarium head blight. Physiol Mol Plant P. 2016;93: 39–48.

20. Buhrow LM, Cram D, Tulpan D, Foroud NA, Loewen MC. Exogenous abscisic acid and gibberellic acid elicit opposing effects on *Fusarium graminearum* infection in wheat. Phytopathology. 2016;106:986–996.

21. Bhattacharya A, Kourmpetli S, Ward DA, Thomas SG, Gong F, Powers SJ, et al. Characterization of the fungal gibberellin desaturase as a 2-oxoglutarate-dependent dioxygenase and its utilization for enhancing plant growth. Plant Physiol. 2012;160:837–45.

22. Luo K, Rocheleau H, Qi P-F, Zheng Y-L, Zhao H-Y, Ouellet T. Indole-3-acetic acid in *Fusarium graminearum*: Identification of biosynthetic pathways and characterization of physiological effects. Fungal Biol. 2016;120:1135–1145.

23. Vrabka J, Niehaus E-M, Münsterkötter M, Proctor RH, Brown DW, Novák O, et al. Production and role of hormones during interaction of Fusarium species with maize (Zea mays L.) seedlings. Front Plant Sci. 2019;9:1936z.

24. Svoboda T, Parich A, Güldener U, Schöfbeck D, Twaruschek K, Václavíková M, et al. Biochemical characterization of the Fusarium graminearum candidate ACC-deaminases and virulence testing of knockout mutant strains. 2019;10:1072.

25. Takeuchi J, Okamoto M, Akiyama T, Muto T, Yajima S, Sue M, et al. Designed abscisic acid analogs as antagonists of PYL-PP2C receptor interactions. Nat Chem Biol. 2014;10:477–82.

26. IWGSC. Shifting the limits in wheat research and breeding using a fully annotated reference genome. Science. 2018;361:eaar7191.

27. Pan Y, Li Y, Liu Z, Surendra A, Wang L, Foroud NA, et al. Differential expression feature extraction (DEFE) and its application in RNA-seq data analysis. bioRxiv. 2019;doi:10.1101/511188.

28. Langfelder P, Horvath S. WGCNA: an R package for weighted correlation network analysis. BMC Bioinformatics. 2008;9:559.

29. Ritchie S, Gilroy S. Abscisic acid signal transduction in the barley aleurone is mediated by phospholipase D activity. Proc Natl Acad Sci U S A. 1998;95:2697–702.

30. Chono M, Matsunaka H, Seki M, Fujita M, Kiribuchi-Otobe C, Oda S, et al. Isolation of a wheat [*Triticum aestivum* L.) mutant in ABA 8⍰-hydroxylase gene: effect of reduced ABA catabolism on germination inhibition under field condition. Breeding Sci. 2013;63:104–115.

31. Prada D, Romagosa I, Ullrich SE, Molina-Cano JL. A centromeric region on chromosome 6(6H) affects dormancy in an induced mutant in barley. J Exp Bot. 2005;56:47–54.

32. Tsai AY, Gazzarrini S. AKIN10 and FUSCA3 interact to control lateral organ development and phase transitions in Arabidopsis. Plant J. 2012;69:809–821.

33. Pan Y, Ouellet T, Phan S, Tchagang A, Fauteux F, Tulpan D. Digitization of trait representation in microarray data analysis of wheat infected by Fusarium graminearum. Proceedings of the 2015 IEEE Conference on Computational Intelligence in Bioinformatics and Computational Biology. 2015;August 12-15, Niagara Falls, Canada.

34. Takino J, Kozaki T, Sato Y, Liu C, Ozaki T, Minami A, et al. Unveiling biosynthesis of the phytohormone abscisic acid in fungi: unprecedented mechanism of core scaffold formation catalyzed by an unusual sesquiterpene synthase. J Am Chem Soc. 2018;140:2392–12395.

35. Becker A, Theissen G. The major clades of MADS-box genes and their role in the development and evolution of flowering plants. Mol Phylogenet Evol. 2003;29:464–89.

36. Cheung AY, Niroomand S, Zou YJ, Wu HM. A transmembrane formin nucleated subapical actin assembly and controls tip-focused growth in pollen tubes. Proc Natl Acad Sci U S A. 2010;107:16390–16395.

37. Bosch M, Helper PK. Pectin methylesterases and pectin dynamics in pollen tubes. Plant Cell. 2005;17:3219–26.

38. Kazan K, Lyons R. Intervention of phytohormone pathways by pathogen effectors. Plant Cell. 2014;26:2285–2309.

39. Foroud NA, Ouellet T, Laroche A, Oosterveen B, Jordan MC, Ellis BE, et al. Differential transcriptome analyses of three wheat genotypes reveal different host response pathways associated with Fusarium head blight and trichothecene resistance. Plant Pathol. 2012;61:296–314.

40. Cuperlovic-Culf M, Rajagopalan N, Tulpan D, Loewen MC. Metabolomics and cheminformatics analysis of antifungal function of plant metabolites. Metabolites. 2016;6:E31.

41. Badea A, Eudes F, Laroche A, Graf R, Doshi K, Amundsen E, et al. Antimicrobial peptides expressed in wheat reduce susceptibility to Fusarium head blight and powdery mildew. Can J Plant Sci. 2013;93:199–208.

42. SeCan (2003) AC Crystal Canada Prairie Spring Red Wheat. SeCan Technical bulletin. 2003. https://secan.com/sites/default/flles/AC%20Crystal.pdf. Accessed 12 Sept 2020.

43. Fathallah-Shaykh HM. Microarrays Applications and Pitfalls. Arch Neuro. 2005;62:1669–1672.

44. Jia H, Cho S, Muehlbauer GJ. Transcriptome analysis of a wheat near-isogenic line pair carrying Fusarium head blight–resistant and -susceptible alleles. Mol Plant Microbe In. 2009;22:1366–1378.

45. Erayman M, Turktas M, Akdogan G, Gurkok T, Inal B, Ishakoglu E, et al. Transcriptome analysis of wheat inoculated with *Fusarium graminearum*. Front. Plant Sci. 2015;6:867.

46. Gunnaiah R, Kushalappa AC, Duggavathi R, Fox S, Somers DJ. Integrated metabolo-proteomic approach to decipher the mechanisms by which wheat QTL (Fhb1) contributes to resistance against *Fusarium graminearum*. Plos One. 2012;7:e40695.

47. Lionetti V, Giancaspro A, Fabri E, Giove SL, Reem N, Zabotina OA, et al. Cell wall traits as potential resources to improve resistance of durum wheat against *Fusarium graminearum*. BMC Plant Biol 2015;15:6.

48. Dhokane D, Karre S, Kushalappa AC, McCartney C. Integrated metabolo-transcriptomics reveals Fusarium head blight candidate resistance genes in wheat QTL-Fhb2. PLoS One. 2016;11: e0155851.

49. Kage U, Karre S, Kushalappa AC, McCartney C. Identification and characterization of a Fusarium head blight resistance gene TaACT in wheat QTL-2DL. Plant Biotechnol J. 2017;15: 447–457.

50. Karre S, Kumar A, Yogendra K, Kage U, Kushalappa A, Charron JB. HvWRKY23 regulates flavonoid glycoside and hydroxycinnamic acid amide biosynthetic genes in barley to combat Fusarium head blight. Plant Mol Biol. 2019;100:591–605.

51. Kang Z, Buchenauer H. Ultrastructural and cytochemical studies on the infection of wheat spikes by *Fusarium culmorum* as well as on degradation of cell wall components and localization of mycotoxins in the host tissue. Mycotoxin Res. 2000;16 Suppl 1:1–5.

52. Mohammadi M, Kazemi H. Changes in peroxidase and polyphenol oxidase activities in susceptible and resistant wheat heads inoculated with *Fusarium graminearum* and induced resistance. Plant Sci. 2002;162:491–498.

53. Siranidou E, Kang Z, Buchenauer H, Studies on symptom development, phenolic compounds and morphological defence responses in wheat cultivars differing in resistance to Fusarium head blight. J Phytopathol. 2003;150, 200–208.

54. Rajagopalan N, Lu Y, Burton IW, Monteil-Rivera F, Halasz A, Reimer E, et al. A phenylpropanoid diglyceride associates with the leaf rust resistance Lr34res gene in wheat. Phytochemistry. 2020;178:112456.

55. Denance N, Sánchez-Vallet A, Goffner D, Molina A. Disease resistance or growth: the role of plant phytohormones in balancing immune responses and fitness costs. Front Plant Sci. 2013;4:155.

56. Long XY, Balcerzak M, Gulden S, Cao W, Fedak G, Wei Y-M, et al. Expression profiling identifies differentially expressed genes associated with the fusarium head blight resistance QTL 2DL from the wheat variety Wuhan-1. Physiol Mol Plant P. 2015;90:1–11.

57. Peng J, Richards DE, Hartley NM, Murphy GP, Devos KM, Flintham JE, et al. Green revolution genes encode mutant gibberellin response modulators. Nature. 1999;400:256–261.

58. Yang W, Yu Z, Yu S, Fan G, Han H, Dong Z, et al. Effect of uniconazole waterless dressing see on yield of wheat. Acta Agron Sin. 2004;30:502–506.

59. Sari E, Cabral AL, Polley B, Tan Y, Hsueh E, Konkin DJ, et al. Weighted gene co-expression network analysis unveils gene networks associated with the Fusarium head blight resistance in tetrapioid wheat. BMC Genomics. 2019;20:925.

60. Buerstmayr M, Lemmens M, Steiner B, Buerstmayr H. Advanced backcross QTL mapping of resistance to Fusarium head blight and plant morphological traits in a *Triticum macha × T. aestivum* population. Theor Appl Genet. 2011;123:293–306.

61. Buerstmayr M, Huber K, Heckmann J, Steiner B, Nelson JC, Buerstmayr H. Mapping of QTL for Fusarium head blight resistance and morphological and developmental traits in three backcross populations derived from *Triticum dicoccum × Triticum durum*. Theor Appl Genet. 2012;125:1751–65.

62. Warth B, Parich A, Bueschl C, Schoefbeck D, Neumann NKN, Kluger B, Schuster K, Krska R, Adam G, Lemmens M, Schuhmacher R. GC–MS based targeted metabolic profiling identifies changes in the wheat metabolome following deoxynivalenol treatment. Metabolomics. 2015;11:722–738.

63. Boedi S, Berger H, Sieber C, Münsterkötter M, Maloku I, Warth B, et al. Comparison of *Fusarium graminearum* transcriptomes on living or dead wheat differentiates substrate-responsive and defense-responsive genes. Front Microbiol. 2016;7:1113.

64. Mihlan M, Homann V, Liu TW, Tudzynski B. AREA directly mediates nitrogen regulation of gibberellin biosynthesis in *Gibberella fujikuroi,* but its activity is not affected by NMR. Mol Microbiol. 2003;47:975–91.

65. Gordon C, Rajagopalan N, Riseeuw E, Surpin M, Ball F, Barber C, et al. Identification of *T. aestivum* abscisic acid receptors and evidence of their role in Fusarium head blight. PLoS One. 2016;11:e0164996.

66. Wang L, He X, Guo J, Shen Y, Huang Z. The expression of wheat TaSTG gene can enhance salt tolerance in plants. Plant Biosyst. 2013;147:451–458.

67. Petsch KA, Mylne J, Botella JR. Cosuppression of eukaryotic Release Factor 1-1 in Arabidopsis affects cell elongation and radial cell division. Plant Physiol. 2005;139: 115–126.

68. Shumayla, Sharma S, Kumar Ry, Mendu V, Singh K, Upadhyay SK. Genomic dissection and expression profiling revealed functional divergence in *Triticum aestivum* leucine rich repeat receptor like kinases (TaLRRKs). Front Plant Sci. 2016;7:I374.

69. Thapa G, Gunupuru LR, Hehir JG, Kahla A, Mullins E, Doohan FM. A pathogen-responsive leucine rich receptor like kinase contributes to Fusarium resistance in cereals. Front Plant Sci. 2018;9:867.

70. Fan X, Naz M, Fan X, Xuan W, Miller AJ, Xu G. Plant nitrate transporters: from gene function to application. J Exp Bot., 2017;68:2463–2475.

71. Krouk G. Hormones and nitrate: a two-way connection. Plant Mol Biol. 2016;91:599–606.

72. Proctore RH, Hohn TM, McCormick SP. Reduced virulence of Gibberella zeae caused by disruption of a tricothecene toxin biosynthetic gene. Mol Plant Microbe In. 8:593–601.

73. Abrams SR, Nelson K and Abrose SJ. Deuterated abscisic acid analogs for mass spectroscopy and metabolism studies. J Labelled Compd Rad. 2003;46:273–283.

74. Lulsdorf MM, Yuan HY, Slater SMH, Vandenberg A, Han X, Zaharia LI, et al. Endogenous hormone profiles during early seed development of *C. arietinum* and *C. anatolicum*. J Plant Growth Regul. 2013;71:191–198.

75. Galka MM, Ambrose SJ, Ross ARS, Abrams SR. Synthesis of deuterated jasmonates for mass spectroscopy and metabolism studies. J. Labelled Compd Rad. 2005;48:797–809.

76. Ross ARS, Ambrose SR, Cutler AJ, Feurtado JA, Kermode AR, Nelson K, et al. Determination of endogenous and supplied deuterated abscisic acid in plant tissues by high performance liquid chromatography-electrospray ionization tandem mass spectroscopy with multiple reaction monitoring. Anal Biochem. 2004;329:324–333.

77. Zaharia LI, Galka MM, Ambrose SJ, Abrams SR. Preparation of deuterated abscisic acid metabolites for use in mass spectroscopy and feeding studies. J Labelled Compd Rad. 2005;48:435–445.

78. Ewing B, Hillier L, Wendl MC, Green P. Base-calling of automated sequencer traces using phred. I. accuracy assessment. Genome Res. 1998;8:175–85.

79. Hannon GJ. FASTX-Toolkit. 2010; http://hannonlab.cshl.edu/fastx_toolkit.

80. Liu Z, Li Y, Pan Y, Wang L, Ouellet T, Fobert P. Strategy in wheat-Fusarium dual-genome RNA-seq data processing. bioRxiv. 2019;doi:10.1101/2019.12.16.878124.

81. Dobin A, Davis CA, Schlesinger F, Drenkow J, Zaleski C, Jha S, et al. STAR: ultrafast universal RNA-seq aligner. Bioinformatics, 2013;29:15–21.

82. Love MI, Huber W, Anders S. Moderated estimation of fold change and dispersion for RNA-seq data with DESeq2. Genome Biol. 2014;15:550.

83. Benjamini Y, Hochberg Y. Controlling the false discovery rate: a practical and powerful approach to multiple testing. J R Stat Soc Ser B Methodol. 1995;57:289–300.

