## Additional File 5 for "Wheat transcriptome profiling reveals abscisic and gibberellic acid treatments regulate early-stage phytohormone defense signaling, cell wall fortification, and metabolic switches following *Fusarium graminearum*-challenge"

### Supplemental Figures

**Transcriptomic and phytohormone profiling reveal early stage molecular variations upon co-application of abscisic or gibberellic acids with *Fusarium graminearum*-challenge of wheat**

**Leann M. Buhrow[1], Ziying Liu[2], Dustin Cram[1], Tanya Sharma[3], Nora A. Foroud[4], Youlian Pan[2]*, Michele C. Loewen[1,3, 5] ***

1. National Research Council of Canada, Aquatic and Crop Resources Development Research Center, 110 Gymnasium Place, Saskatoon, SK, S7N 0M8, Canada
2. National Research Council of Canada, Digital Technologies Research Center, 1200 Montreal Road, Ottawa, ON, K1A 0R6, Canada
3. University of Ottawa, Department of Chemistry, 150 Louis-Pasteur Pvt, Ottawa, ON, K1N 6N5, Canada
4. Agriculture and Agri-food Canada, Lethbridge Research and Development Center, 5403 1st Ave, Lethbridge, AB, T1J 4B1, Canada
5. National Research Council of Canada, Aquatic and Crop Resources Development Research Center, 100 Sussex Drive, Ottawa, ON, K1A 0R6, Canada

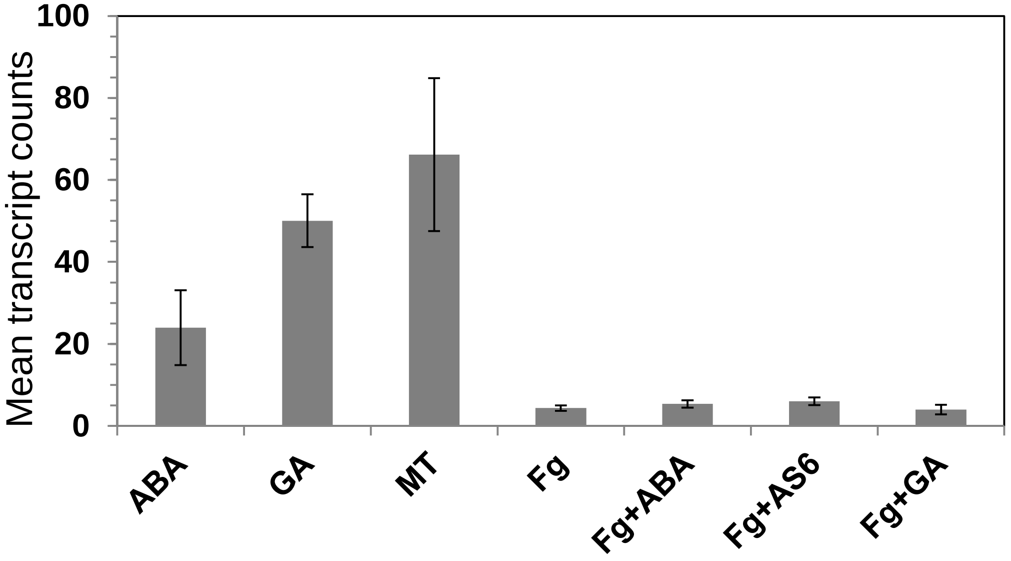


**Supplemental Figure S1.** Abundance of wheat FUS3 transcript (TraesCS3D01G249100) under various treatments. Error bars represent standard error of the mean.

**Supplemental Figure S2:** *Fusarium graminearum* transcriptomic data quantification. **A)** Number of *Fg* DEGs of each pairwise comparison. **B)** Expression profile for FGRAMPH1_01T26277 transcripts, with homology to the *B. cinera* ABA biosynthetic gene, BcCPR1. Error bars represent standard error of the mean.

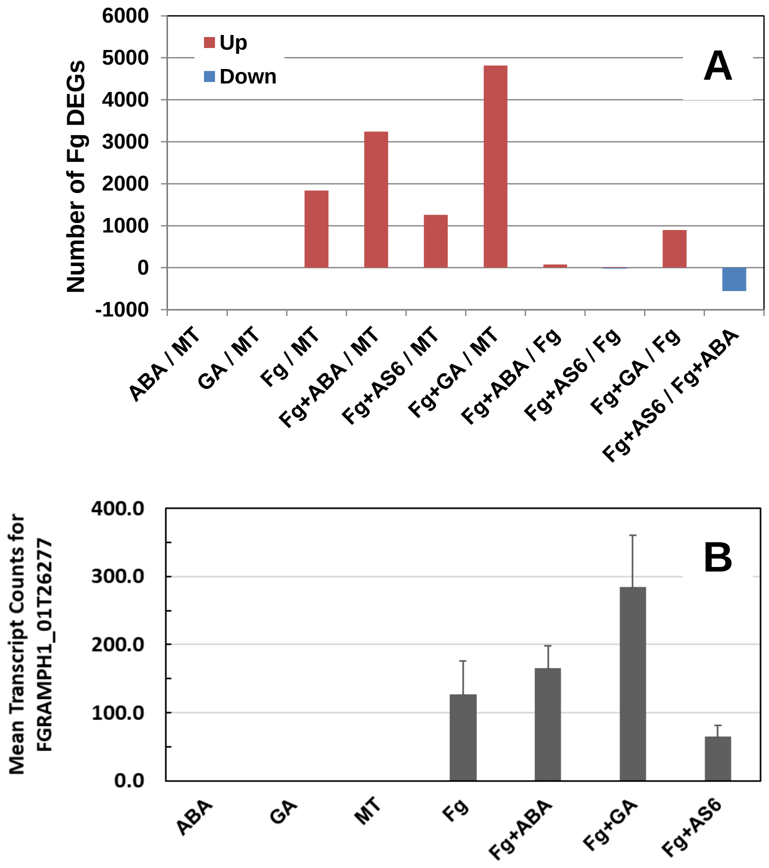

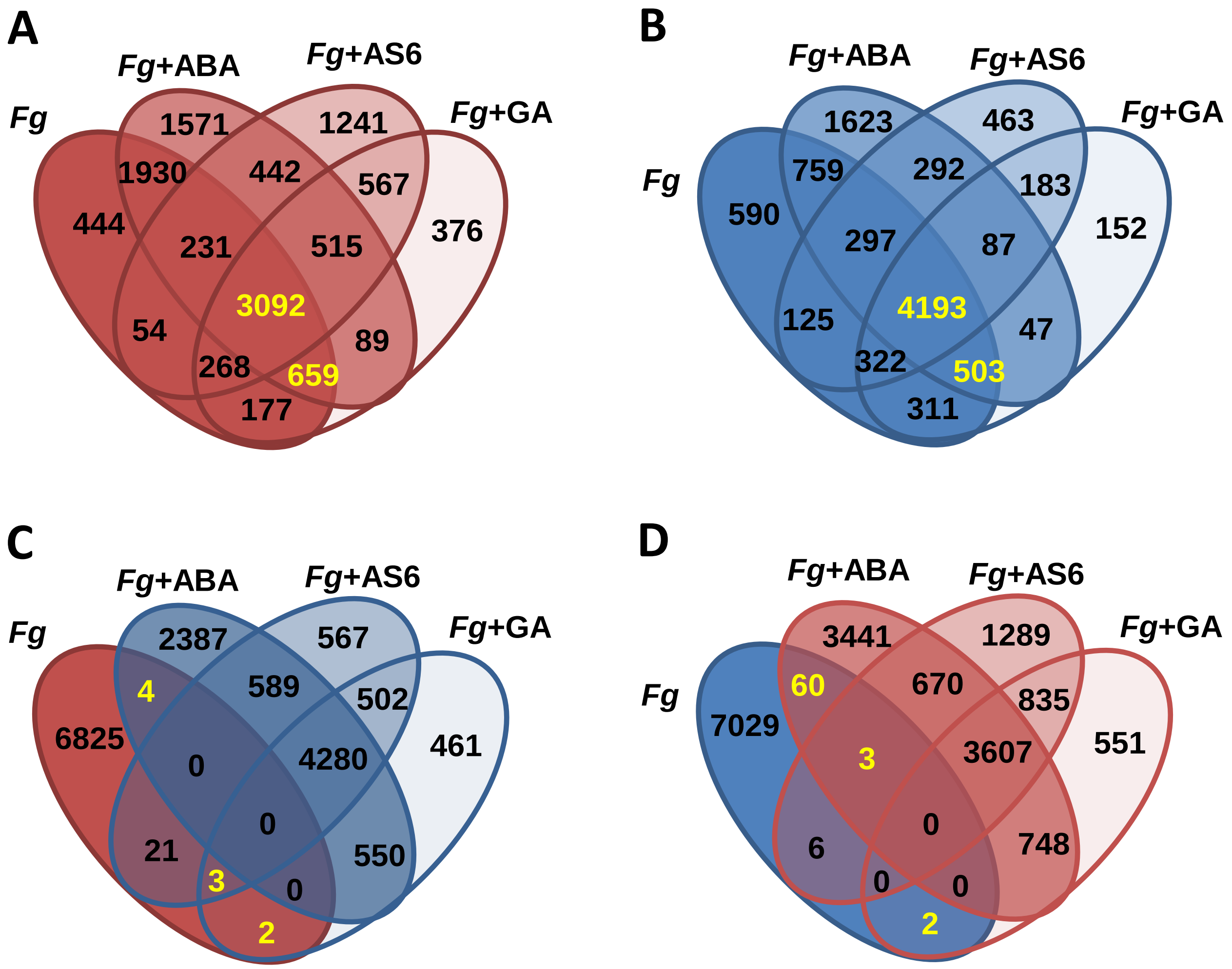


**Supplemental Figure S3. Share of *Fg* alone up- or down- regulated genes that are similarly up- and down- regulated under alternate conditions, with all DEGs calculated compared to mock treated (MT).** In all instances, upregulated genes are highlighted in shades of red, and downregulated genes are highlighted in shades of blue. DEGs called in text are yellow. **A)** Among the 6855 genes up-regulated by *Fg* that were also up-regulated under the co-applied conditions of *Fg*+ABA or *Fg*+GA or *Fg*+AS6. **B)** Among the 7100 genes down-regulated by *Fg* that were also down-regulated under the co-applied conditions of *Fg*+ABA or *Fg*+GA or *Fg*+AS6. **C)** Up-regulated by *Fg*, but down-regulated under the co-applied conditions of *Fg*+ABA or *Fg*+GA or *Fg*+AS6. **D)** Down-regulated by *Fg*, but up-regulated under the co-applied conditions of *Fg*+ABA or *Fg*+GA or *Fg*+AS6.

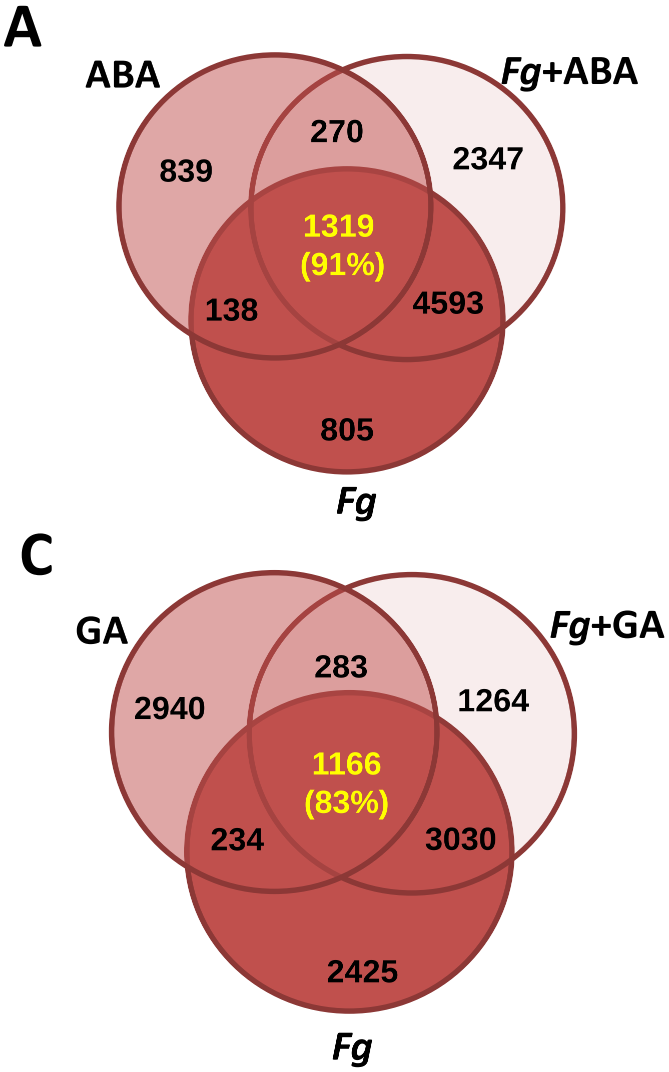


**Supplemental Figure S4. Comparison of gene regulation elicited by ABA and GA in the presence and absence of *Fg*, with DEGs calculated compared to mock treated (MT) 'Fielder'.** In all cases, upregulated genes are highlighted in shades of red, and downregulated genes are highlighted in shades of blue. DEGs called in text are yellow. **A)** Distribution of all upregulated genes by ABA alone or *Fg*+ABA and *Fg*-alone. **B)** Distribution of all down-regulated genes by ABA alone, *Fg*+ABA and *Fg*-alone. **C)** Distribution of all upregulated genes by GA alone or *Fg*+GA and *Fg*-alone. D) Distribution of all down-regulated genes by GA alone, *Fg*+GA and *Fg*-alone.

**A**

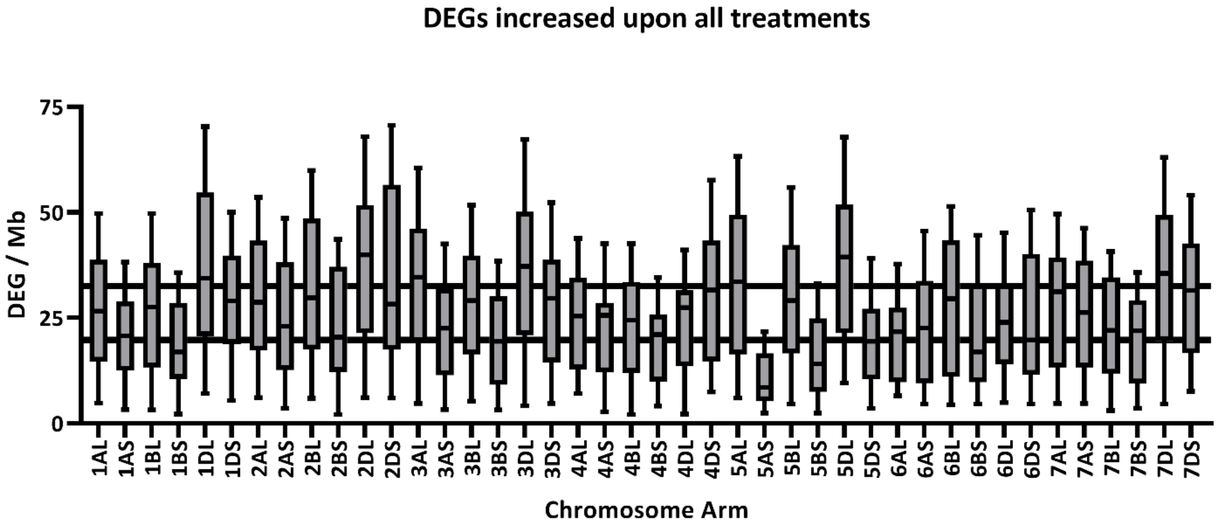


**B**

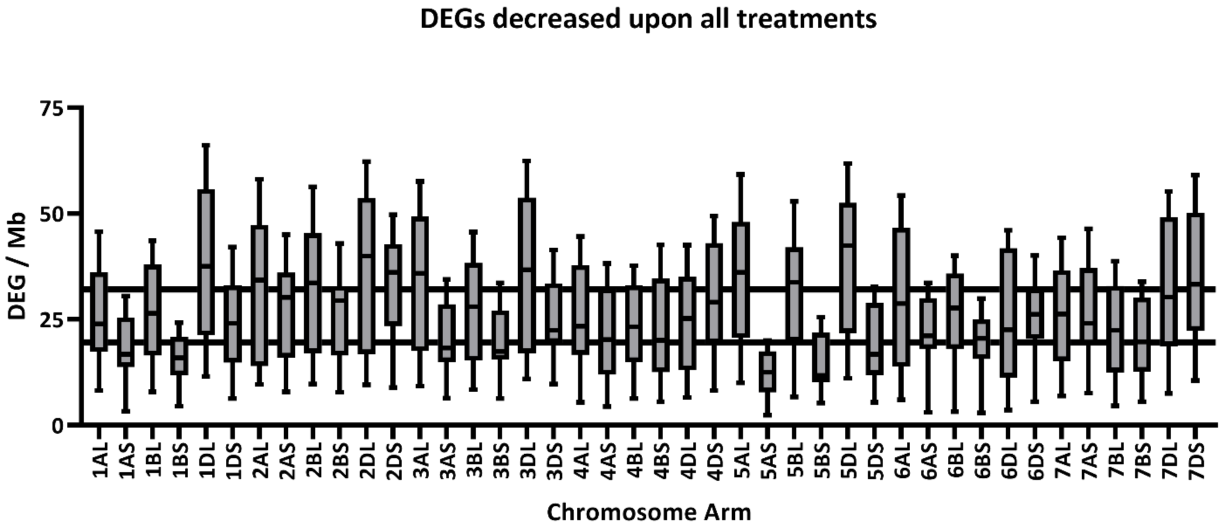


**Supplemental Figure S5: Chromosomal density of mapped wheat DEGs elicited by *Fg* and phytohormone treatments.** Density was calculated as mapped wheat DEG count induced upon ABA, GA, *Fg*, *Fg*+ABA, *Fg*+GA, and *Fg*+AS6 treatments divided by physical chromosome arm length (physical lengths were derived from IWGSC, 2018). DEG density was relatively monodisperse where chromosome arms had a global average of A) 26.1 ± 6.4 up-regulated and B) 25.9 ± 6.3 down-regulated DEGs. The variation in these global averages ± 1 standard deviation are represented by lateral lines.

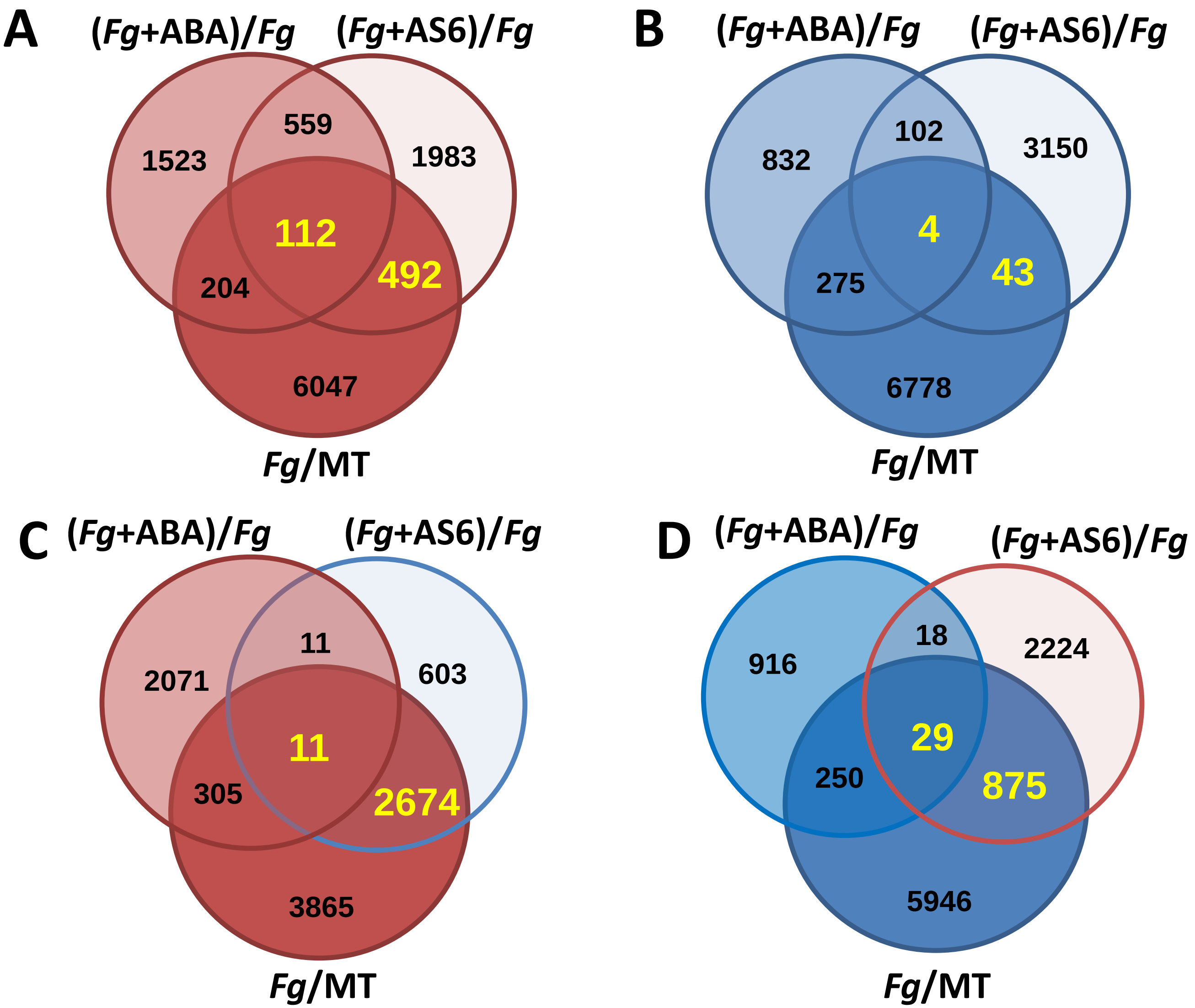


**Supplemental Figure S6. Comparison of enhancement and suppression ABA and AS6 in the presence of *Fg* as compared to *Fg* alone.** In all cases, upregulated genes are highlighted in shades of red, and downregulated genes are highlighted in shades of blue. DEGs called in text are yellow. **A)** Distribution of all upregulated genes by *Fg*+ABA and *Fg*+AS6 as compared to *Fg*, or *Fg*-alone compared to MT. **B)** Distribution of all down-regulated genes by *Fg*+ABA and *Fg*+AS6 as compared to *Fg*, or *Fg*-alone compared to MT. **C)** Distribution of up-regulated genes by *Fg*-alone, further enhanced by *Fg*+ABA , but inhibited by *Fg*+AS6 as they compared to *Fg*. **D)** Distribution of down-regulated genes by *Fg*-alone, further enhanced by *Fg*+ABA, but inhibited by *Fg*+AS6 as they compared to *Fg*.

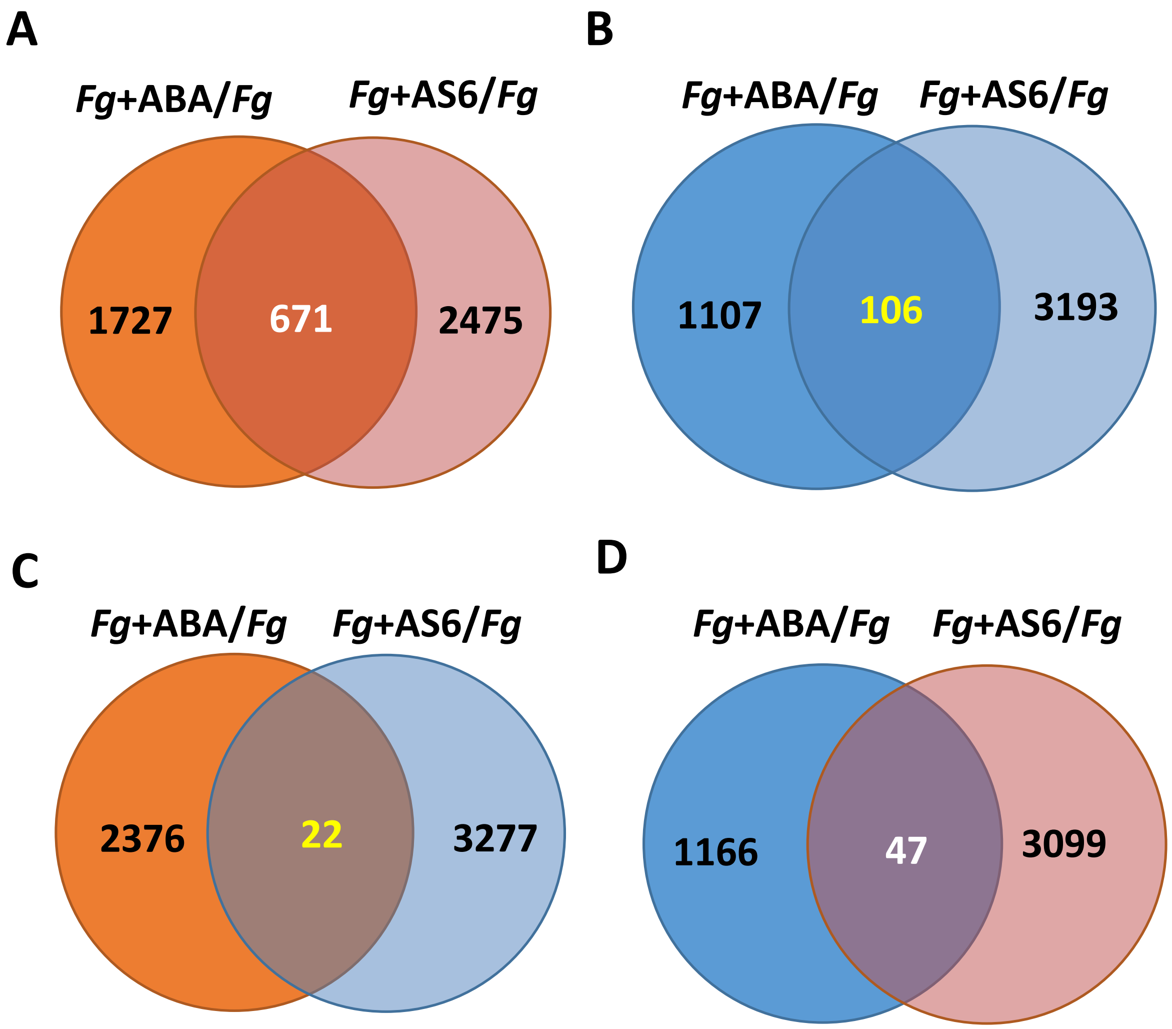


**Figure S7: Share of enhanced or suppressed genes by co-applied ABA and AS6, with all DEGs calculated and compared to *Fg*-alone.** In all instances, up-regulated genes are highlighted in shades of red, and down-regulated genes are highlighted in shades of blue. **A)** All up-regulated DEGS. **B)** All down-regulated DEGS. Opposing DEGs: **C)** up-regulated by *Fg*+ABA but downregulated by *Fg*+AS6; **D)** down-regulated by *Fg*+ABA but upregulated by *Fg*+AS6. Enhanced genes are highlighted in white, suppressed genes are highlighted in yellow.

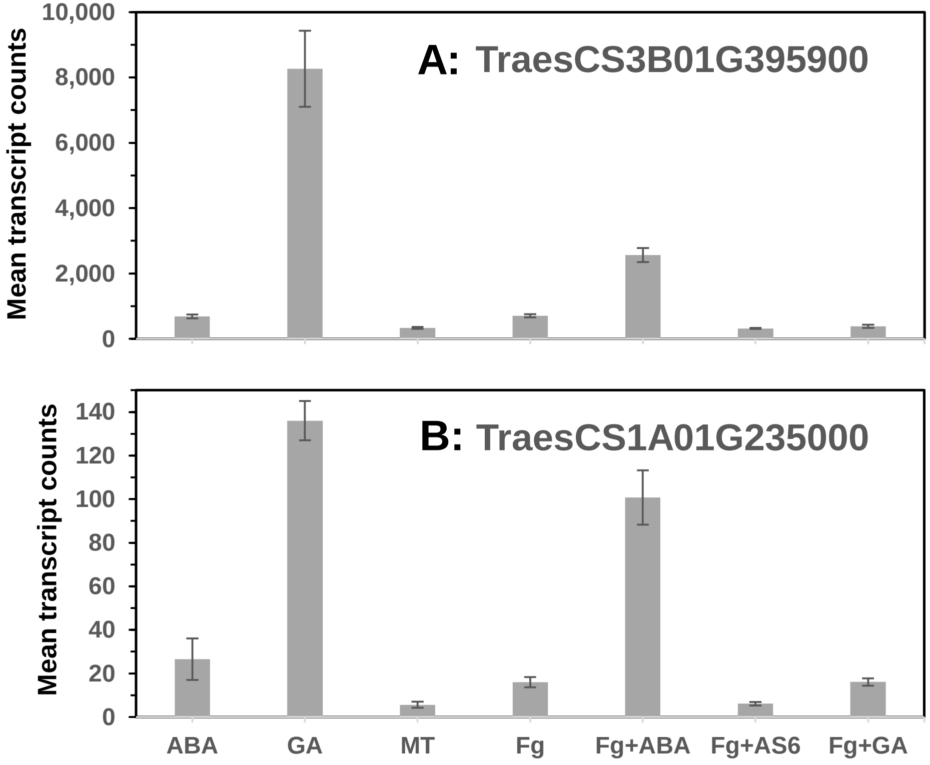


**Supplemental Figure S8. Expression profile of two genes up-regulated by ABA with or without *Fg*, and by *Fg* alone, but down-regulated by *Fg*+AS6. A)** a gamma-glutamyl phosphate reductase. **B)** a eukaryotic peptide chain release factor subunit 1-1. Error bars represent standard error of the mean.

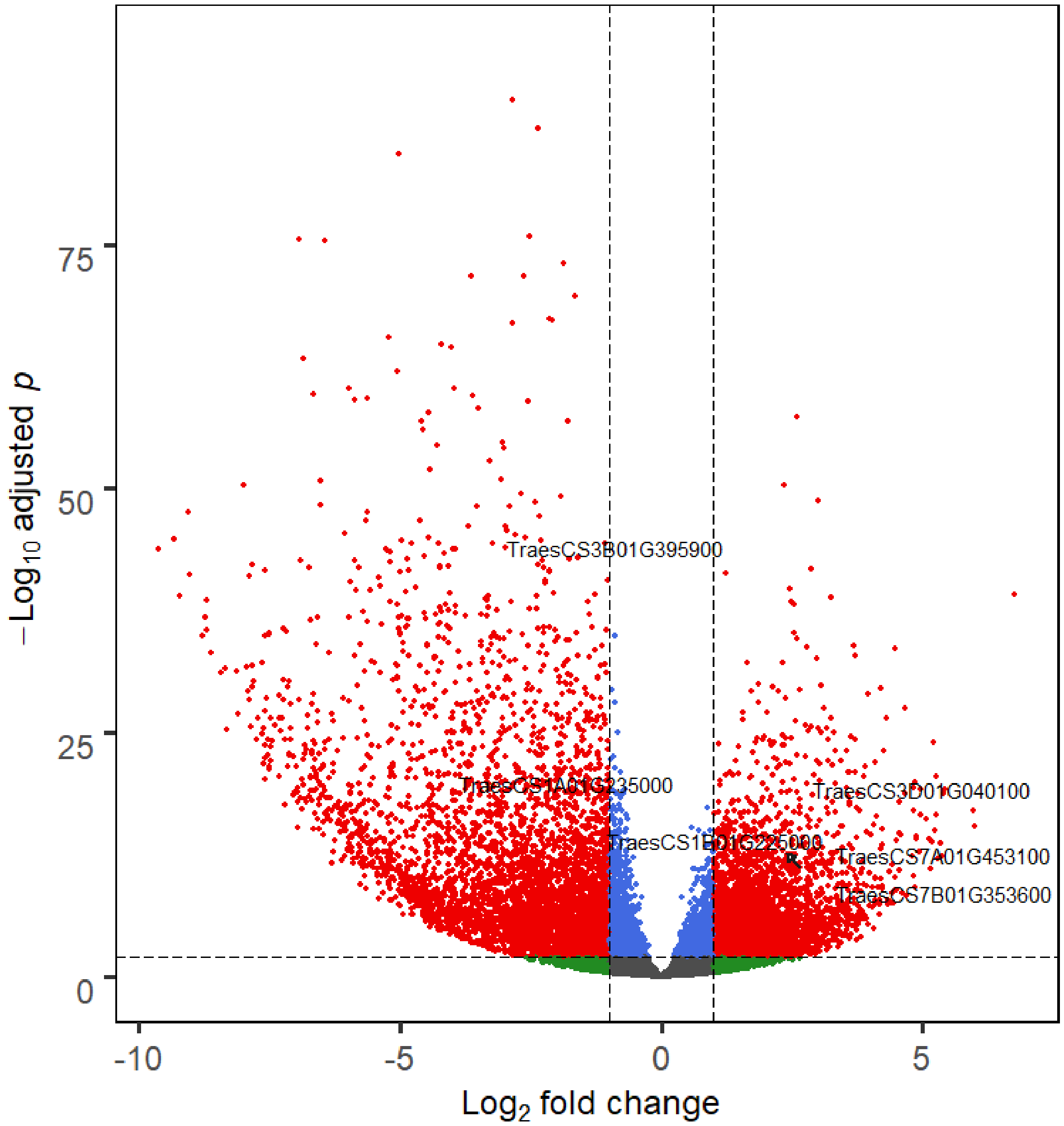


**Supplemental Figure S9. Differentially expressed genes between conditions of *Fg*+AS6 and *Fg*+ABA.** Their difference in the six genes (Figures S8 and S10) are all highly significant (adj $p < 1E\text{-}10$).

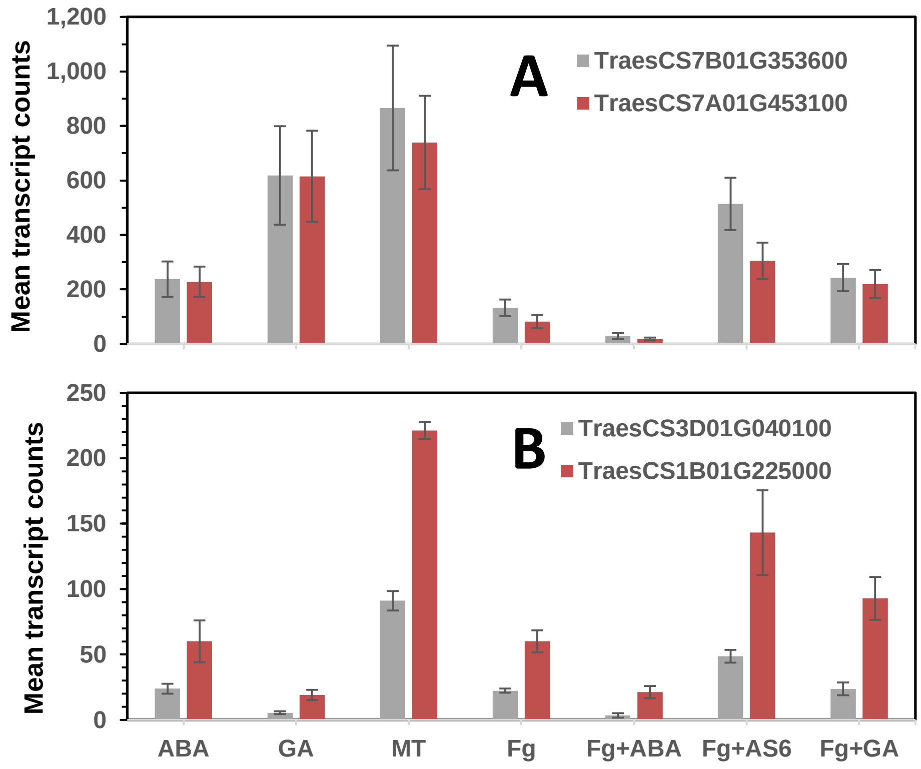


**Supplemental Figure S10. Expression profile of four genes down-regulated by ABA with or without *Fg*, and by *Fg* alone, but such down-regulation were mitigated by *Fg*+AS6. A)** two transmembrane proteins. **B)** a Leucine-rich repeat receptor-like protein kinase family protein and a Nitrate transporter 1.1. Error bars represent standard error of the mean.

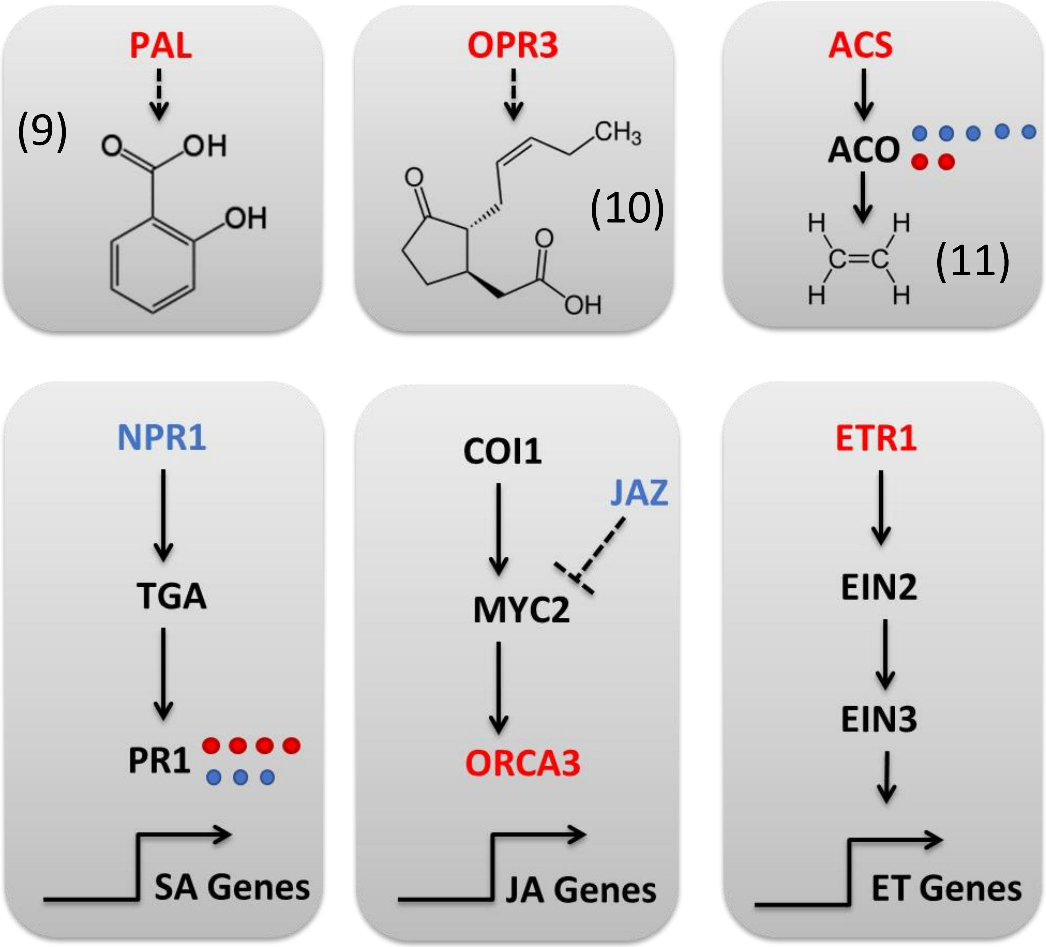


**Supplemental Figure S11: *Fg* challenge alters the expression of 'Fielder' classical defense hormone biosynthesis and signaling pathway genes.** 'Fielder' gene acronyms are represented as upregulated (red; log2FC > 1.5) and down regulated (blue; log2FC < -1.5), respectively (adj p < 0.01) compared to mock treatment. Annotations were based on Blast analysis selecting for transcripts with > 50 % sequence identity to known phytohormone signaling pathway members as annotated in the KEGG database. Chemical structures are shown for (9) salicylic acid, (10) jasmonic acid and (11) ethylene. Detailed expression data is located in **Additional file 1, Tabs S2** and **S3**.

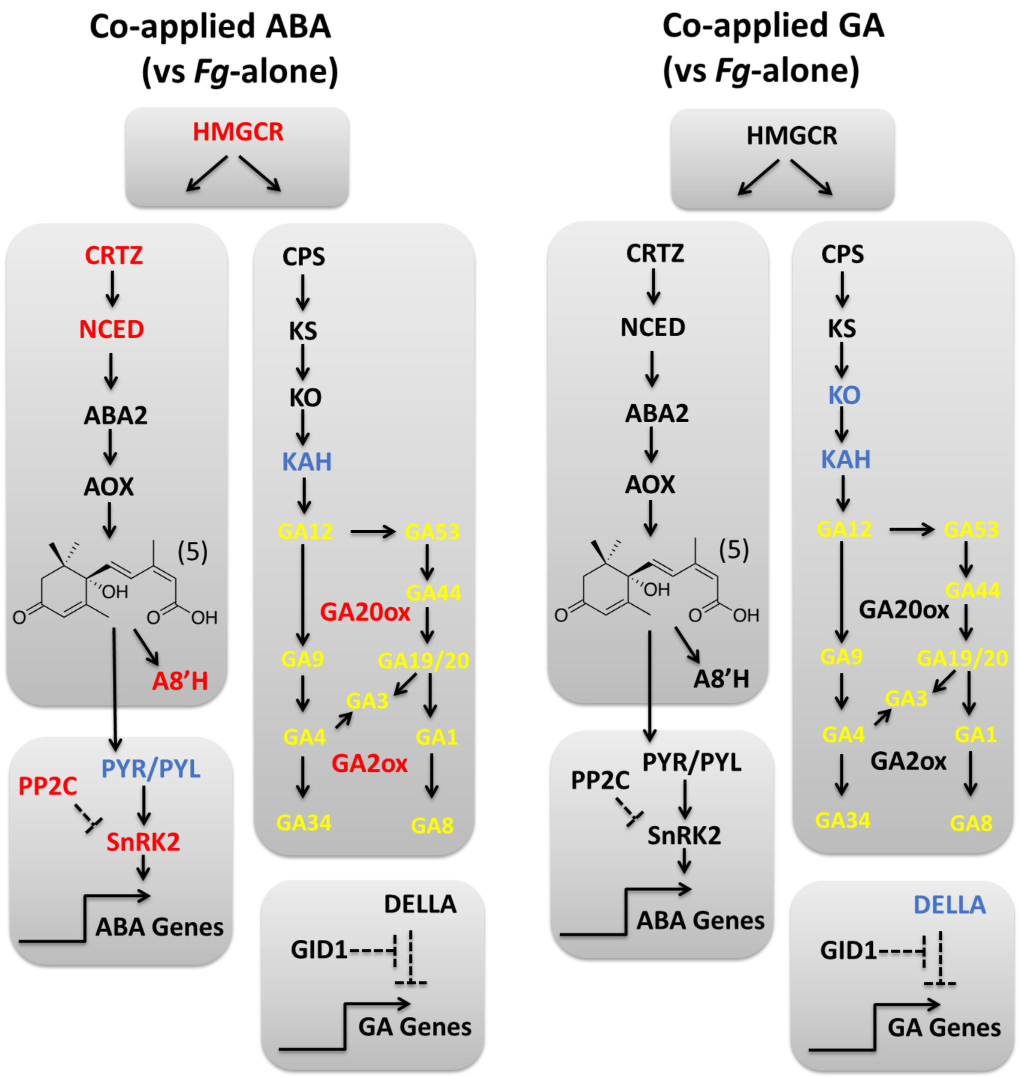


**Supplemental Figure S12: The co-application of ABA and GA with *Fg* leads to modulation of their respective pathways, with DEGs calculated compared to *Fg* alone.** Chemical structures are shown for (5) abscisic acid. GA metabolite acronyms are highlighted in yellow and the rest of the legend details are the same as in **Figure S11**. Detailed expression data is located in **Additional file 1, Tabs S2** and **S3**.

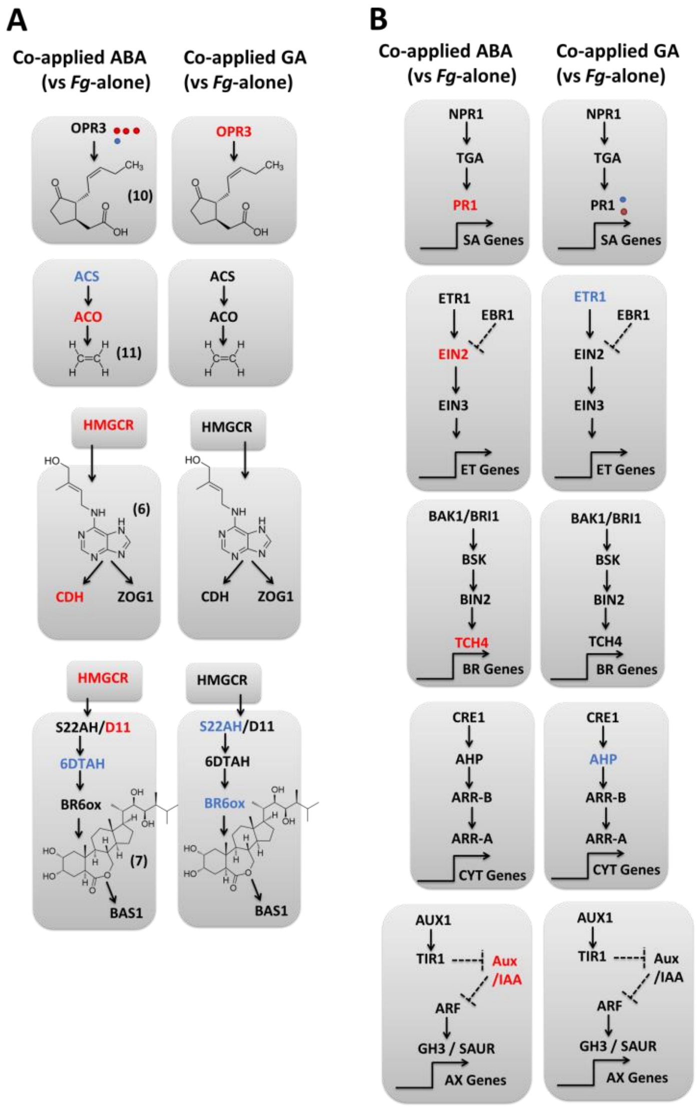


**Supplemental Figure S13: Phytohormone biosynthetic and signaling responses arising from co-application of ABA and GA with *Fg* challenge, with DEGs calculated compared to *Fg*-alone. A)** Biosynthetic pathways with chemical structures for (10) jasmonic acid, (11) ethylene, (6) trans-zeatin an (7) brassinolide, see **Additional file 1, Tab S2** for details. **B)** Signaling pathways, see **Additional file 1, Tab S3** for details. Legend details are otherwise the same as in **Figure S11**.
